# Proximity labeling at H3K9me3 reveals VRK-1 regulate global chromatin distribution in *C. elegans*

**DOI:** 10.64898/2026.07.07.737140

**Authors:** William Smith, Valeryia Aksianiuk, Ramon Pfaendler, Rodrigo Villaseñor, Devanarayanan Siva Sankar, Michael Stumpe, Peter Lenart, Peter Askjaer, Benjamin D. Towbin, Tuncay Baubec, Jörn Dengjel, Peter Meister

## Abstract

Heterochromatin marked by histone H3 lysine 9 di- or trimethylation (H3K9me2/3) underpins transcriptional silencing and nuclear organization, yet its full complement of associated proteins remains incompletely defined. Here, we apply ChromID proximity labelling with the mouse HP1β chromodomains to map the H3K9me3-proximal proteome in *Caenorhabditis elegans*, recovering known heterochromatin factors alongside previously uncharacterized candidates. We pursued one such candidate, the vaccinia-related kinase VRK-1, because of its established but poorly understood links to chromatin organization. Intriguingly, VRK-1 dynamically relocates from a broad nuclear distribution to the nuclear periphery upon azide or heat stress. Following these stresses, bulk chromatin exhibits similarly increased peripheral enrichment and apparent compaction, as assessed by radial fluorescence profiles. Although VRK-1 is not necessary for stress-induced chromatin reorganization, decompaction and repositioning of chromatin away from the nuclear envelope during recovery requires VRK-1. VRK-1 depletion leads to persistent perinuclear chromatin retention and compromises post-stress survival. Furthermore, loss of VRK-1 catalytic activity results in over-retention of chromatin at the nuclear periphery under normal growth conditions; this phenotype can be reversed by depletion of a key VRK-1 substrate at the nuclear envelope BAF-1. Our findings identify VRK-1 as a key regulator that controls the interaction of chromatin with the nuclear lamina through regulation of BAF-1.

## Introduction

Eukaryotic genomes are organized into transcriptionally active euchromatin and compact, gene-poor transcriptionally more silent heterochromatin (Heitz 1928; Frenster et al. 1963). A defining hallmark of constitutive heterochromatin is trimethylation of histone H3 on lysine 9 (H3K9me3), deposited by the Suv39h-family methyltransferases and recognized by Heterochromatin Proteins 1 (HP1-family chromodomain proteins) (Djeghloul et al. 2023). These factors contribute to chromatin compaction, although the degree of compaction is dynamic and modulated by processes such as DNA repair and replication (Dialynas et al. 2007; Strom et al. 2017). In the vast majority of organisms and cell types, H3K9me3-rich domains are enriched at the nuclear periphery or cluster into chromocentres; however, this organization varies considerably (Solovei et al. 2009; Politz et al. 2013).

In the nematode *Caenorhabditis elegans*, chromatin organization follows these principles but with unique features; *C. elegans* has holocentric chromosomes, so classic pericentromeric heterochromatin blocks are absent (Mandrioli and Manicardi 2020). Instead, H3K9 methylation marks numerous small repressive domains enriched on the distal arms of each chromosome (Ikegami et al. 2010; Towbin et al. 2012; Liu et al. 2011). This distribution correlates with genetic elements: H3K9 methylation targets repetitive elements such as transposons and simple repeats, as well as a subset of tissue-inappropriate genes, while its absence leads to upregulation of repetitive elements (Zeller et al. 2016). A wide array of factors recognizes H3K9 methylation to target, establish, maintain and localize heterochromatin within the nucleus.

Methyltransferases MET-2 and SET-25 deposit H3K9me1/2 and H3K9me1/2/3, respectively, primarily in a stepwise manner (Towbin et al. 2012). To deposit H3K9me3, SET-25 is recruited to preexisting H3K9me2 deposited by MET-2, through a direct interaction with the MBT-domain protein LIN-61, which recognizes the dimethylated mark (Padeken et al. 2021). The small RNA interference (RNAi) pathway independently promotes SET-25 recruitment through the argonaut protein NRDE-3 (Padeken et al. 2021). The highly conserved heterochromatin binding protein (HP1 or CBX1 in humans and mice) is a well characterised reader of H3K9me2/3 (Schoelz and Riddle 2022). Their *C. elegans* orthologues HPL-1/-2, similarly recognize methylated H3K9 and regulate chromatin compaction and gene expression (Schott et al. 2006; Garrigues et al. 2015; de la Cruz-Ruiz et al. 2023), while chromodomain protein CEC-4 tethers H3K9me-marked chromatin to the inner nuclear membrane (Gonzalez-Sandoval et al. 2015). Beyond these proteins, the full repertoire of proteins that associate with H3K9me3-marked chromatin in *C. elegans* remains largely undefined or uncharacterised.

To address this gap, we employed ChromID, a proximity-labeling strategy in which an engineered chromodomain reader module fused to a promiscuous biotin ligase biotinylates proteins near specific histone modifications, here H3K9me3 (Villaseñor et al. 2020). Because the biotin ligase labels proteins within an approximately 10 nm radius, ChromID captures both stably bound factors as well as freely diffusing proteins that transiently interact with H3K9me3, hence providing an unbiased approach to probe the protein composition at a given histone mark.

Applying ChromID with the mouse CBX1 chromodomain in *C. elegans* L3 larvae, we identified the vaccinia-related kinase VRK-1 as an H3K9me3-proximal factor. VRK-1 is a conserved nuclear Ser/Thr kinase with well-established roles in chromosome biology. A central function of VRK family kinases across species is the phosphorylation of BAF-1 (barrier to autointegration factor), a small DNA-bridging protein that mediates interactions between chromatin and the nuclear lamina. Phosphorylation of BAF-1 by VRK-1 reduces its affinity for both DNA and inner nuclear membrane proteins, and this regulation is essential for post-mitotic nuclear envelope reassembly in multiple systems (for reviews see(Lazo 2024; Campillo-Marcos et al. 2021)). Beyond mitosis, BAF-1 mobility at the nuclear periphery is regulated by VRK-1 during heat stress in *C. elegans* (Bar et al. 2014), and recent work has shown that VRK-1 drives detachment of meiotic chromosomes from the nuclear periphery through BAF-1 in the *C. elegans* germline (Paouneskou et al. 2025). Despite these connections to chromatin-lamina interactions, VRK-1 had not been directly linked to heterochromatin regulation or identified as a component of the H3K9me3-proximal proteome.

Here, we show that VRK-1 redistributes to the nuclear periphery upon heat or azide stress, and surprisingly these stresses also induce global chromatin compaction and perinuclear accumulation. In both cases these effects are reversible during recovery. Critically, chromatin decompaction and release from the nuclear envelope during recovery require VRK-1. Furthermore, in catalytically inactive *vrk-1* mutants, chromatin is constitutively compacted and enriched at the nuclear periphery even under non-stress conditions. Importantly, this is a phenotype largely suppressed by depletion of BAF-1. Although VRK-1 was identified through its proximity to H3K9me3, our functional data indicate that it acts as a broad regulator of chromatin-lamina interactions through BAF-1, rather than functioning specifically at heterochromatin/H3K9me3. Together, these findings reveal a conserved VRK-1-BAF-1 signaling axis that controls global chromatin distribution during stress and normal growth.

## Results

### Mouse Cbx1/HP1-β chromodomain dimers bind specifically to nematode trimethylated Histone 3 Lysine 9

To identify the protein composition at trimethylated H3K9 in *C. elegans*, we used ChromID, a proximity labeling approach originally established in mouse ESCs(Villaseñor et al. 2020). To establish the specificity of the trimethyl-H3K9 binder in *C. elegans,* two copies of the CBX1 chromodomain (CBX1_2x) were fused to an NLS and mScarlet and expressed from a single-copy transgene driven by the ubiquitous *dpy-30* promoter (Fig. 1A). CBX1_2x formed a speckled nuclear pattern enriched at the nuclear periphery (Fig. 1B), consistent with known H3K9me3 distribution (Towbin et al. 2012; Ikegami et al. 2010). CBX1_2x colocalized with condensed chromatin domains labeled by histone H3.3 HIS-72::GFP and with a well-characterized heterochromatic reporter array marked by H3K9me2/3 and H3K27me3 (Fig. 1C)(Meister et al. 2010; Towbin et al. 2012). To confirm specificity, we expressed the CBX1_2x::mScarlet in a *set-25* mutant that lacks detectable H3K9me3 while retaining H3K9me2 (Towbin et al. 2012). In this mutant background, the perinuclear enrichment and foci was abolished and CBX1_2x::mScarlet became diffuse throughout the nuclear lumen (Fig. 1D, radial profiling in 1E-F (Padovani et al. 2022)). These results demonstrate that CBX1_2x chromodomains bind specifically to H3K9me3 heterochromatin in *C. elegans*.

**Figure 1.**
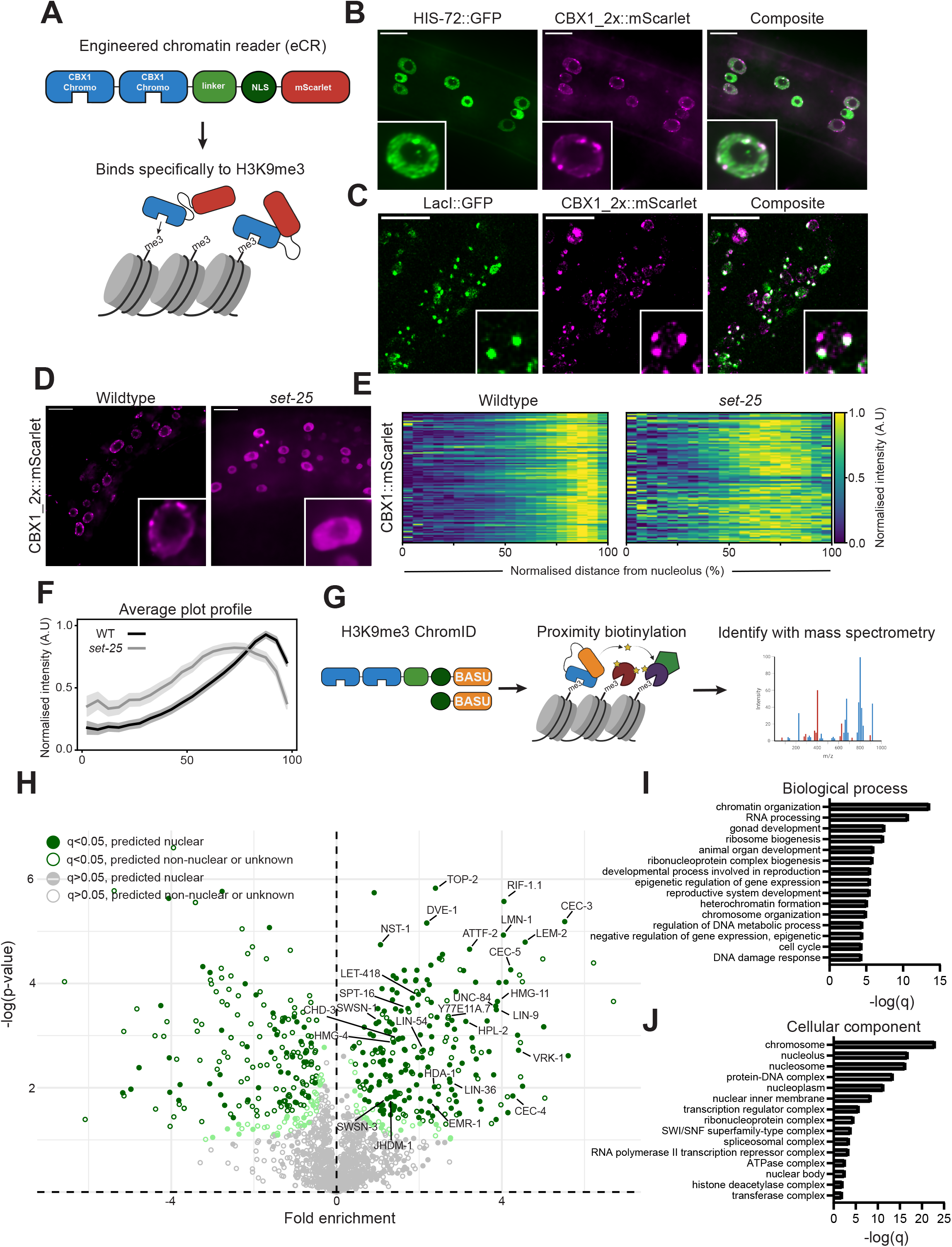
CBX1_2x proximity labelling identifies the protein composition at H3K9me3 in *C. elegans*. **A.** Schematic diagram of the H3K9me3 engineered chromatin reader. Two copies of the CBX1 chromodomain (CBX1_2x) were fused to an NLS and mScarlet. The construct was inserted as a single copy using MosSCI and is driven under a ubiquitous *dpy-30* promoter. **B.** Live confocal images of CBX1_2x::mScarlet and HIS-72::GFP in hypodermal cells, where animals were immobilised with levamisole. Scale bars indicate 10 μM **C.** CBX1_2x::mScarlet colocalises with *gwIs4*, a highly repetitive, H3K9me2/3 and H3K27me3 heterochromatic array anchored at the nuclear periphery (about 300 plasmid repeats). *gwIs4* contains *lacO* sites bound by GFP-lacI and can be visualised *in vivo*. Scale bar is 2 μm **D.** Representative single plane images of CBX1_2x::mScarlet in wildtype and *set-25(n5021)* mutant hypodermal nuclei. *set-25* mutants lack detectable H3K9me3. Scale bars indicate 10 μM **E.** Heatmaps displaying the average normalised nuclear distribution of CBX1_2x::mScarlet in wildtype and *set-25(n5021)* hypodermal nuclei from the nucleolus (left) to the nuclear periphery (right) (n= 111, 91 for wildtype and *set-25* respectively). Normalised signal intensity is binned into 5% intervals. Individual rows correspond to single nuclei. **F.** Average plot profiles for the nuclear distribution of CBX1_2x::mScarlet in wildtype and *set-25* nuclei, as described and measured in E. Dark lines indicate the average, whereas shaded regions indicate 95% confidence intervals (Al-Refaie et al. 2024). **G.** Schematic diagram of CBX1_2x::BASU ChromID. The CBX1_2x chromodomain was fused to the promiscuous biotin ligase BASU and inserted as a single copy using MosSCI. CBX1_2x::BASU biotinylates proteins in an approximately 10 nm radius around CBX-1_2X bound H3K9me3 histones. High abundance endogenously biotinylated mitochondrial carboxylases were His10 tagged and depleted from the protein extract. Biotinylated proteins were isolated using streptavidin pulldown and identified using LC-MS/MS. The fold changes are log2 transformed and calculated relative to free NLS::BASU, to control for background nuclear biotinylation. Approximately 100,000 worms were grown in parallel, in triplicate for CBX1_2x::BASU and NLS::BASU, and harvested at the L3 larval stage. **H.** Volcano plot displaying significantly enriched proteins proximal to CBX1_2x::BASU relative to NLS::BASU in L3 worms. Fold changes are log2 transformed and calculated relative to NLS::BASU to control for background nuclear biotinylation. Significantly enriched proteins with q<0.05 are labeled dark green and significantly enriched proteins with 0.05≤q<0.1 are labeled light green. Detected proteins were categorised as predicted nuclear, indicated by filled circle, or predicted non-nuclear/unknown, indicated with an open circle using the cellular component gene ontology from UniProtKB entries aligned to protein IDs. Green indicates significantly enriched hits (q<0.05). Ribosomal and mitochondrial proteins were censored using the cellular component ontology. After filtering, 251 proteins were identified as significantly enriched in CBX1_2x::BASU against NLS::BASU. **I.** Bar chart showing the top 15 most significantly enriched biological process (BP) terms of the 251 positively enriched hits (PANTHER 19.0). Bars show -log (q) values. Generic terms (>500 genes associated) and overspecific terms (<10 genes associated) were filtered out. Redundant terms were manually filtered. **J.** As above, but using cellular component ontology (CC). Generic terms were also removed, as described above.

### ChromID identifies the protein composition at H3K9me3

We next fused CBX1_2x chromodomains to the promiscuous biotin ligase BASU to perform proximity labeling (Fig. 1G). Nematodes expressing CBX1_2x::BASU or the control untargeted NLS::BASU were grown to the third larval stage (L3), and biotinylated proteins were isolated by streptavidin pulldown after biochemical depletion of highly abundant endogenously biotinylated mitochondrial carboxylase (Artan et al. 2022). Liquid chromatography-tandem mass spectrometry (LC-MS/MS) in triplicate identified 1864 proteins, of which 251 proteins were significantly enriched over the NLS::BASU control (q<0.1), with 220 of high confidence (q<0.05) (Fig. 1H; supplementary table 1).

Among the enriched proteins were established H3K9me3-associated proteins: HPL-2, which binds H3K9me3 *in vivo* (Gonzalez-Sandoval et al. 2015; de la Cruz-Ruiz et al. 2023); CEC-4, the nuclear envelope tether for H3K9me3 (Gonzalez-Sandoval et al. 2015; de la Cruz-Ruiz et al. 2023)); CEC-3/EAP-1 and CEC-5, which recognize H3K9me1/2/3 *in vitro*, and colocalize *in vivo* with H3K9me2/3 for CEC-3 or associate strongly with H3K9me2 for CEC-5 (Greer et al. 2014; Hou et al. 2023); and inner nuclear membrane components LMN-1 (lamin), EMR-1, LEM-2 (LEM-domain protein), and UNC-84 (SUN domain protein) (Bizhanova and Kaufman 2021; Ahringer and Gasser 2018). Notably however, we did not detect the H3K9 methyltransferase SET-25, its recruitment factor LIN-61, or the second HP1 homolog HPL-1 across all pulldowns (Padeken et al. 2019; Meister et al. 2011; Towbin et al. 2012). Additional enriched factors included the histone deacetylase HDA-1, implicated in piRNA-mediated silencing (Kim et al. 2021a); the putative H3K9me2 histone demethylase JHDM-1(Lee et al. 2019); and components of FACT (Facilitates and Contrasts Transcription), NuRD (nucleosome remodelling and deacetylation), SynMuvB/DRM, and SWI/SNF complexes (Passannante et al. 2010; Suggs et al. 2018; Rechtsteiner et al. 2019; Hoareau et al. 2024; Large and Mathies 2014). Among proteins not previously linked to H3K9me3 in *C. elegans*, were the high mobility group proteins HMG-11/12, the vaccinia-related kinase VRK-1 and the uncharacterized disordered and unstudied protein Y77E11A.7, which shares structural homology with SETDB1.

Gene ontology (GO) enrichment analysis of the H3K9me3-proximal proteome highlighted biological process terms including heterochromatin formation, epigenetic regulation of gene expression, and RNA processing (Fig. 1I), and cellular component terms including nucleolus, nuclear inner membrane, and transcription regulator complex (Fig. 1J). Altogether, ChromID identifies known H3K9me3-associated factors and subnuclear compartment residents, validating the approach, while revealing an array of factors with diverse nuclear functions not previously linked to H3K9me3. We focused further on VRK-1, given that loss of VRK-1 has been reported to increase H3K9me3 levels in mammalian cells (Monte-Serrano et al. 2023) and to cause visibly condensed chromatin in *C. elegans* and zebrafish (Gorjánácz et al. 2007; Carrasco Apolinario et al. 2023).

### VRK-1 and chromatin relocate to the nuclear periphery upon azide stress

To investigate VRK-1 function at trimethylated H3K9 heterochromatin, we characterized its localization in somatic nuclei using an endogenously tagged allele (*bq28*, *vrk-1::wrmScarlet*), together with GFP-labelled histone H3.3 (*his-72*) (Delaney et al. 2018). When animals were immobilized with levamisole, an agonist of nicotinic acetylcholine receptors leading to muscle contraction (Davis and Tanis 2022), VRK-1 in hypodermal cells was broadly distributed in the nuclear space, with reduced intensity in the nucleolus and colocalized with a subset of chromatin foci (Fig. 2A, levamisole), consistent with interphase VRK1 distribution observed in mammalian cells (Monte-Serrano et al. 2023). Notably, VRK-1 distribution did not recapitulate the focal, peripherally enriched pattern of CBX1_2x::mScarlet (Fig. 1B), indicating that VRK-1 is not restricted to H3K9me3-marked heterochromatin but is broadly associated with chromatin across the nucleus.

**Figure 2.**
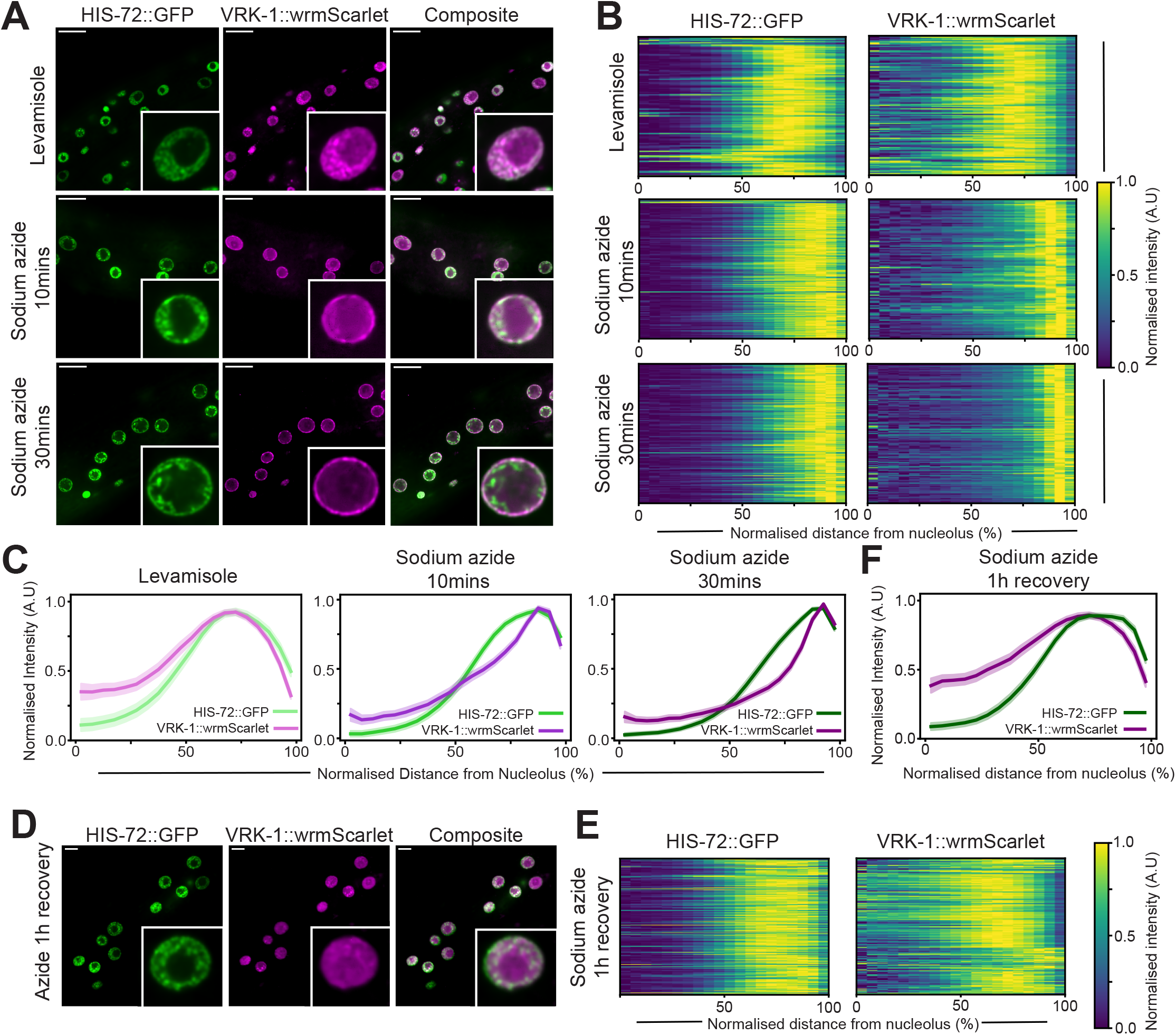
Sodium azide but not levamisole treatment results in the reversible relocation of VRK-1 and HIS-72::GFP to the nuclear periphery. **A.** Single plane representative images of chromatin, labelled with HIS-72::GFP and VRK-1::wrmScarlet in L3/4 hypodermal nuclei. Animals were imaged after 20-30 mins on 2% agarose pads containing either 10 mM levamisole or 0.1% sodium azide at indicated timepoints (10 mins, 30 mins). Scale bar 10 μm **B.** Heatmaps displaying normalised intensity from nucleolus/nuclear centre to nuclear periphery for indicated immobiliser or fluorophore, where each row is an individual nucleus. n=127, n=124 for levamisole, HIS-72::GFP, VRK-1::wrmScarlet, respectively. n=170, n=118 for sodium azide 10 mins HIS-72::GFP, VRK-1::wrmScarlet, respectively. n=191, n=160 for sodium azide 30 mins HIS-72::GFP, VRK-1::wrmScarlet, respectively. **C.** Average plot profiles of all nuclei for indicated immobilized and timepoint. Colour scheme as in 1F. **D.** Representative confocal images of HIS-72::GFP and VRK-1::wrmScarlet after 0.1% sodium azide treatment in M9 for 30 minutes and 1h recovery. Scale bars indicate 5 μm. For images/quantifications of M9 treatment after 30minutes no recovery or levamisole see supplementary figure S1. **E.** Heatmaps showing normalised signal intensity for images in D., as described in B. **F**. Average plot profiles for sodium azide recovery as described above. n>235 for HIS-72::GFP and VRK-1::wrmScarlet. **F.** Average plot profiles from quantifications made in E.

Strikingly, immobilization with sodium azide triggered rapid redistribution of VRK-1 from a diffuse irregular nucleoplasmic pattern to the nuclear periphery (Fig. 2A, sodium azide). This perinuclear enrichment increased over time (Fig. 2A, sodium azide, compare 10 with 30 minutes), indicating progressive VRK-1 relocation (Fig. 2A). In parallel, chromatin accumulated at the nuclear rim and appeared more condensed over time, although this relocation was less rapid and pronounced than for VRK-1 (20-30 minutes, Fig. 2B, HIS-72, sodium azide). The differential kinetics of VRK-1 and chromatin redistribution argue both against a simple model where chromatin bound VRK-1 relocated alongside chromatin and in which chromatin and VRK-1 relocation is passively displaced by nucleolar expansion, although we cannot exclude a contribution from changes in nucleolar volume. These changes occurred in postmitotic cells, ruling out cell-cycle effects or relocation due to the mitotic function of VRK-1 (Sulston and Horvitz 1977). Radial fluorescence profiling confirmed the relocation of both VRK-1 and chromatin towards the nuclear periphery in hypodermal cells (Fig. 2BC) (Padovani et al. 2022).

Next, we assessed whether the effects of sodium azide on the nuclear distribution of VRK-1 and chromatin were reversible. In our standard imaging protocol, animals are exposed to sodium azide directly on agarose pads, which does not allow the drug to be washed out; we therefore performed levamisole/sodium azide treatment in liquid M9 buffer instead. This treatment produced the same relocation of chromatin and VRK-1 within 30 minutes (Fig. S1A-C). When animals were subsequently allowed to recover for one hour, the distribution of both chromatin and VRK-1 largely returned to that of untreated animals (Fig. 2DE), although VRK-1 appears somewhat more diffuse within the nucleus than under untreated conditions. Similar results were obtained for intestinal nuclei (Fig. S1D-F), although chromatin relocation was less pronounced, likely because intestinal nuclei have a large nucleolus and chromatin is already predominantly perinuclear under control conditions.

The delay between the rapid accumulation of VRK-1 at the nuclear rim and the slower relocation of chromatin to the same location suggests that VRK-1 is sequestered at the nuclear envelope either independently of, or only partially together with chromatin.

Together, our experiments demonstrate that sodium azide, but not levamisole, drives the rapid but reversible relocation of VRK-1 towards the nuclear periphery in multiple tissues. Sodium azide also has a strong effect on genome architecture in hypodermal cells, causing compaction and a shift towards the nuclear periphery that is likewise largely reversible. We therefore used levamisol as the immobilization agent for all subsequent experiments unless otherwise stated, and focused our analyses on hypodermal nuclei.

### Heat shock induces relocalisation of VRK-1 and chromatin to the nuclear rim

To determine whether the relocation of VRK-1 and chromatin represents a component of stress response, we examined the effects of heat shock. Animals were imaged during heat shock (after 30 min at 37°C) as well as during short-term (30 min) and long-term (24 h) recovery following a 1 hour heat shock at 37°C (Fig. 3A). As for sodium azide, VRK-1 became enriched at the nuclear periphery during heat shock, although to a somewhat lesser extent than with azide (Fig. 3BC, heatmaps Fig. S2). Concurrently, chromatin accumulated at the nuclear periphery and the nucleoplasmic fraction appeared highly condensed (Fig. 3BC, heatmaps Fig. S2). During early recovery (30 min), chromatin distribution largely resembled that during heat shock, whereas VRK-1 redistributed broadly across chromatin, colocalizing extensively with dense chromatin foci (Fig. 3B). After 24 h, chromatin distribution was largely restored in most nuclei, though it remained slightly more condensed than in unstressed conditions (Fig. 3BC, heatmaps Fig. S2). Notably, VRK-1 remained peripherally enriched in approximately half of the nuclei at 24 h (Fig. 3BC, heatmaps in S2).

**Figure 3.**
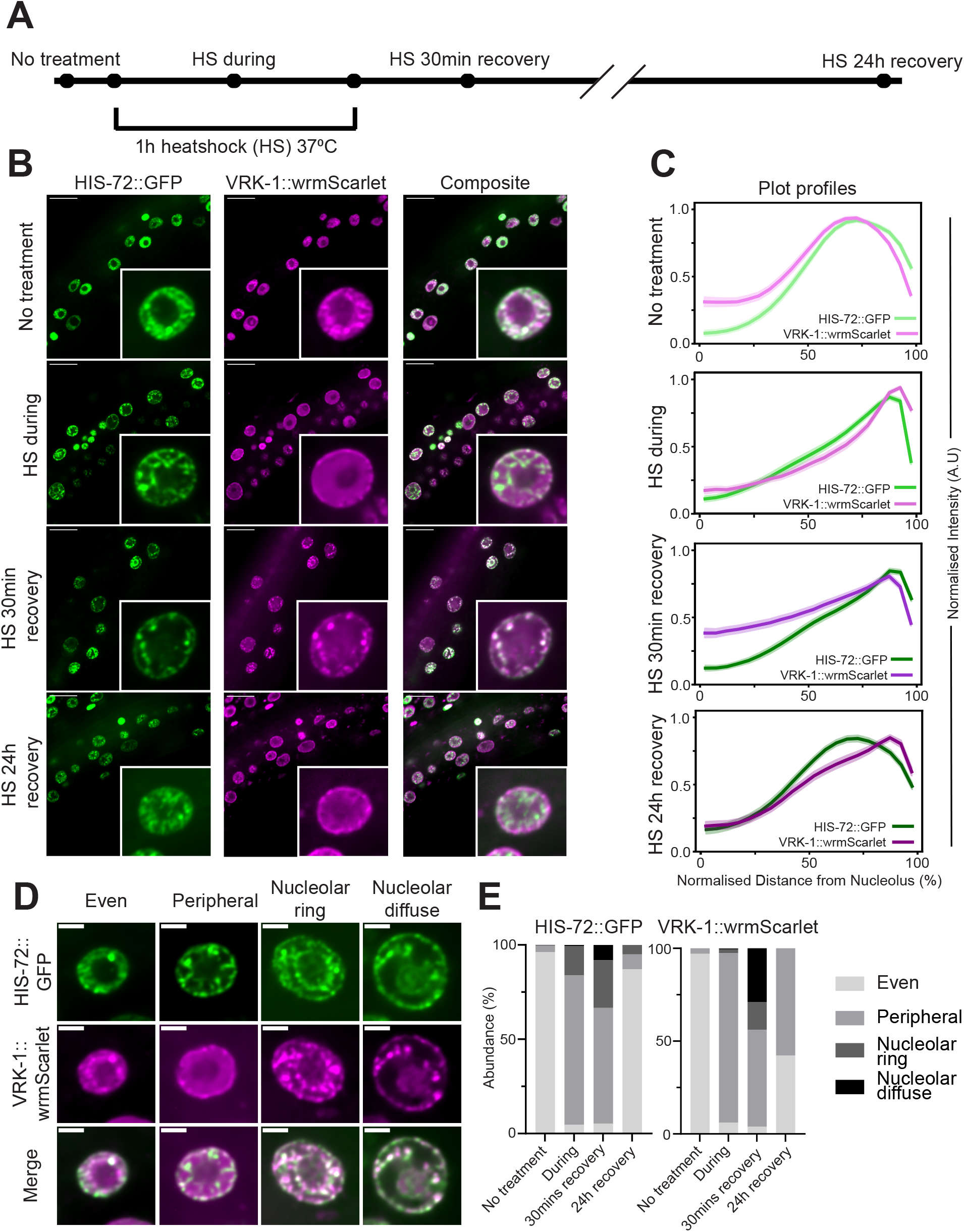
Heat shock results in the reversible relocation of VRK-1::wrmScarlet and HIS-72::GFP to the nuclear periphery in hypodermal cells. **A.** Timeline showing imaging time points before and after 1 hour heat shock at 37°C. Late stage larvae (L3/L4) were imaged on 2% agarose pads containing 3 mM levamisole. For the time point “HS during” nematodes were incubated in a microscope chamber set at 37°C and imaged after 30 minutes. For recovery timepoints, animals were heat shocked on a PCR block at 37°C in 60 mm NGM plates (12 ml of NGM) and imaged 30 minutes or 24 hours after heat shock. **B.** Representative single plane confocal images of HIS-72::GFP and VRK-1::wrmScarlet in hypodermal nuclei before heat shock, during, and after 30 minutes or 24h recovery, as described in A. Scale bar indicates 10 μm. **C.** Average nuclear plot profiles for HIS-72::GFP and VRK-1::wrmScarlet of normalised signal intensity from nucleolus/nuclear centre to nuclear periphery at indicated timepoints, as described in 1f. (n=408, 516, 569, 304 for no treatment, *in situ*, 30 mins recovery and 24h recovery, for HIS-72::GFP, respectively) (n= 408, 516, 569, 304 for no treatment, in situ, 30 mins recovery and 24h recovery for VRK-1::wrmScarlet, respectively). **D.** Representative nuclei of different nuclear distribution categories of HIS-72::GFP and VRK-1::wrmScarlett induced by heat shock. Scale bar 2 μm. Nuclei were categorised by manual inspection of image and its individual plot profile from C. (see methods) **E.** Bar charts showing abundance (%) of VRK-1::wrmScarlet and HIS-72::GFP nuclear conformations observed from heat shock. n numbers are the same as for C.

We further assessed whether our observations from heat shock or sodium azide treatment reflected bulk chromatin changes, and not redistribution of this specific histone variant (H3; HIS-72::GFP), by imaging a different histone variant, H2B (mCherry::HIS-11). We obtained comparable results, indicating that the observed relocation induced by heat shock and sodium azide reflects bulk chromatin redistribution (Fig. S3A–C).

However, we observed significant heterogeneity in chromatin and VRK-1 distributions following heat-shock, particularly regarding enrichment around or within the nucleolus during short-term recovery (Fig. S2). We therefore scored nuclei into four categories; uniform distribution throughout the nucleoplasm (even), peripheral, nucleolar ring, and nucleolar diffuse (Fig. 3D) (see Methods). During heat shock, 95% of nuclei showed altered chromatin distribution, with the majority (79%) exhibiting perinuclear enrichment (Fig. 3E). During early recovery, the proportion of nuclei with nucleolar enrichment increased to 20% compared to 12% during heat stress. Importantly, 87% of nuclei returned to an even chromatin distribution after 24h of recovery. For VRK-1, 57% of nuclei retained perinuclear enrichment after 24 hours recovery (Fig. 3E). Overall, these data demonstrate that, similar to sodium azide, heat shock induces large scale, reversible reorganization of nuclear architecture, with VRK-1 dynamically redistributing between chromatin and the nuclear periphery during recovery. Although the two stresses elicit similar changes, their dynamics differ. Most notably, heat shock requires a substantially longer recovery period.

### VRK-1 is crucial for chromatin reorganization during stress recovery

Given that VRK-1 is a chromatin-associated kinase that relocates to the nuclear periphery during stress, and previous literature suggest that chromatin is over-retained at the nuclear periphery in the absence of VRK-1 during cell division (Lancaster et al. 2007; Paouneskou et al. 2025), we hypothesized that VRK-1 contributes to chromatin recovery following stress. We generated animals carrying an auxin-inducible degron tagged VRK-1::wrmScarlet, in a background that ubiquitously expresses the TIR1 E3 ligase in somatic tissues (Zhang et al. 2015). Auxin (IAA) treatment from the first larval stage led to efficient VRK-1 degradation, with no detectable fluorescence signal in third larval stage animals (Fig. 4A).

**Figure 4.**
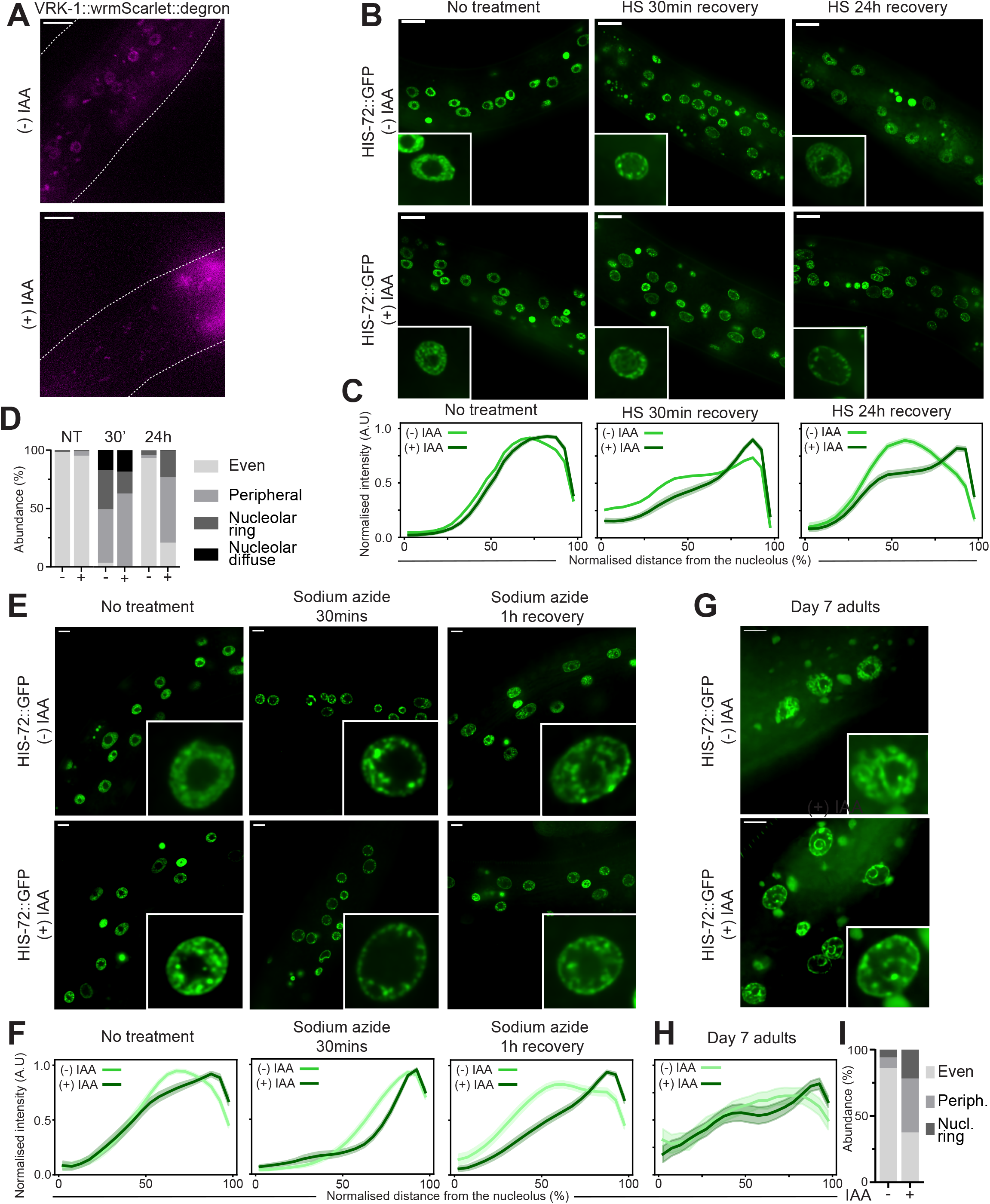
VRK-1 is required for the proper recovery of chromatin distribution after heat shock or sodium azide treatment, and its long-term depletion results in similar ectopic peripheral chromatin. **A.** Single plane confocal representative images of VRK-1::mScaret::degron in the presence or absence of IAA. Dotted lines indicate animal outline, asterisks intestinal autofluorescence. Scale bar 10 μm. **B.** Single plane representative images of HIS-72::GFP in L3 hypodermal nuclei. Scale bar 5 μm. Animals were synchronised using a 2-3 hours egglay on media containing ethanol or 1 mM IAA, grown for 48 hours at 20°C, heat shocked for 1 hour at 37°C and imaged before HS, after 30 minutes recovery or 24h recovery, as described in figure 3. **C.** Average plot profiles for (-) or (+) IAA at timepoints indicated in B. and calculated as described in 1F. (n=580, 257, 167 for (-) IAA no treatment, 30 mins recovery and 24h recovery, respectively. (n= 228, 350, 301 for (+) IAA no treatment, 30 mins recovery and 24h recovery, respectively). **D.** Bar charts displaying percent (%) abundance of nuclear signal distribution for heat shock time points (NT=no treatment, 30’=30 minute recovery and 24h=24h recovery, as described in Fig. 3DE. **E.** Representative single plane confocal images of HIS-72::GFP after VRK-1 depletion ((+) IAA or control ((-) IAA) for sodium azide azide recovery experiment as described in Fig. 2/Fig. S1. Animals were exposed to 0.1% sodium azide in M9 buffer. (n>235 for HIS-72::GFP and VRK-1::wrmScarlet). Scale bar: 5 μm. **F.** Average plot profiles for quantifications of images in E. at indicated timepoints, measured as described in Fig. 1. **G.** Single plane representative images of hypodermal nuclei in day 7 adults having grown with or without IAA, scale bar 5 μm. **H.** Average plot profiles from quantifications of radial signal distribution of nuclei in images from G. (n=50, n=64 (-)/(+) IAA, respectively). **I.** (%) abundance of nuclear distribution categories -/+ IAA in day 7 adults, measured as described in Fig. 3DE.

In contrast to homozygous *vrk-1* null mutations, somatic depletion of VRK-1 over several generations did not produce protruding vulva defects (Klerkx et al. 2009) (data not shown), likely because incomplete knockdown left sufficient VRK-1 to retain partial function. Under control conditions, VRK-1 depletion caused a subtle increase in perinuclear chromatin at the L3 stage (Fig. 4BC, EF, S4AB). During early recovery from heat shock (30 min), VRK-1-depleted animals displayed increased perinuclear chromatin enrichment relative to controls, with variable nucleolar enrichment (Fig. 4BD, S4A). Importantly, 24 hours post-heat shock, 79% of nuclei in VRK-1-depleted animals retained compact perinuclear chromatin, whereas control nuclei had largely recovered (Fig. 4BD).

We next tested whether chromatin relocation/recovery induced by sodium azide would behave in a similar manner to heat shock after VRK-1 depletion. In this case, chromatin became substantially more enriched at the nuclear periphery during azide treatment in the absence of VRK-1 (Fig. 4EF, S4B). Importantly, similar to heat shock, normal chromatin distribution failed to fully recover after one hour in VRK-1 depleted animals (Fig. 4EF, S4B). Notably however, chromatin is slightly less enriched at the nuclear envelope after 1h recovery compared to during sodium azide treatment after VRK-1 depletion. Therefore, it is possible even in the absence of VRK-1, chromatin is still able to recover, albeit dramatically slower than the control.

These data suggest that VRK-1 is critical for chromatin redistribution away from the nuclear periphery during stress recovery. Because we had observed a small increase in peripheral chromatin following VRK-1 depletion under non-stress conditions, we next asked whether VRK-1 acts continuously to prevent perinuclear chromatin accumulation in normal conditions. We therefore depleted VRK-1 for an extended period (seven days) in adult animals using auxin. This led to a dramatic increase in perinuclear and perinucleolar relocation and compaction of chromatin (Fig. 4G-I, S4C). RNAi-mediated *vrk-1* knockdown produced comparable results (Fig. S4DE). This suggests depletion of VRK-1 leads to a progressive accumulation of chromatin at the nuclear periphery over time. Altogether, we conclude that VRK-1 is required both for proper chromatin recovery after stress and for preventing its accumulation at the nuclear rim under normal conditions.

### Loss of the catalytic activity of VRK-1 results in the perinuclear compaction of the genome

VRK1 is a serine/threonine kinase with multiple characterized targets in mammalian cells, including transcription factors ATF2 and p53 as well as the nuclear lamina protein BAF1 (barrier to autointegration factor 1) (Lazo 2024). To determine whether the kinase activity of VRK-1 is required for its chromatin-regulatory function, we complemented the *vrk-1(ok1181)* null mutant with a kinase-dead (KD) allele, *vrk-1(K169E)::mCherry*, expressed from a single-copy transgene under the *vrk-1* promoter, as described previously (Dobrzynska and Askjaer 2016). As reported, first-generation *vrk-1^KD^* homozygotes derived from heterozygous mothers are viable owing to the maternal contribution of wild-type VRK-1 protein, but are sterile because VRK-1 is absent from their germline (Dobrzynska and Askjaer 2016). In animals lacking catalytically active VRK-1, chromatin was highly enriched at the nuclear periphery in 71% of hypodermal nuclei in L4 larvae under normal growth conditions (Fig. 5A-C, S5A). We note this is a significantly stronger effect compared to depletion of VRK-1 by AID degron, where we previously found only a mild increase in peripheral chromatin under non-stress conditions in late larvae (Fig. 4)

**Figure 5.**
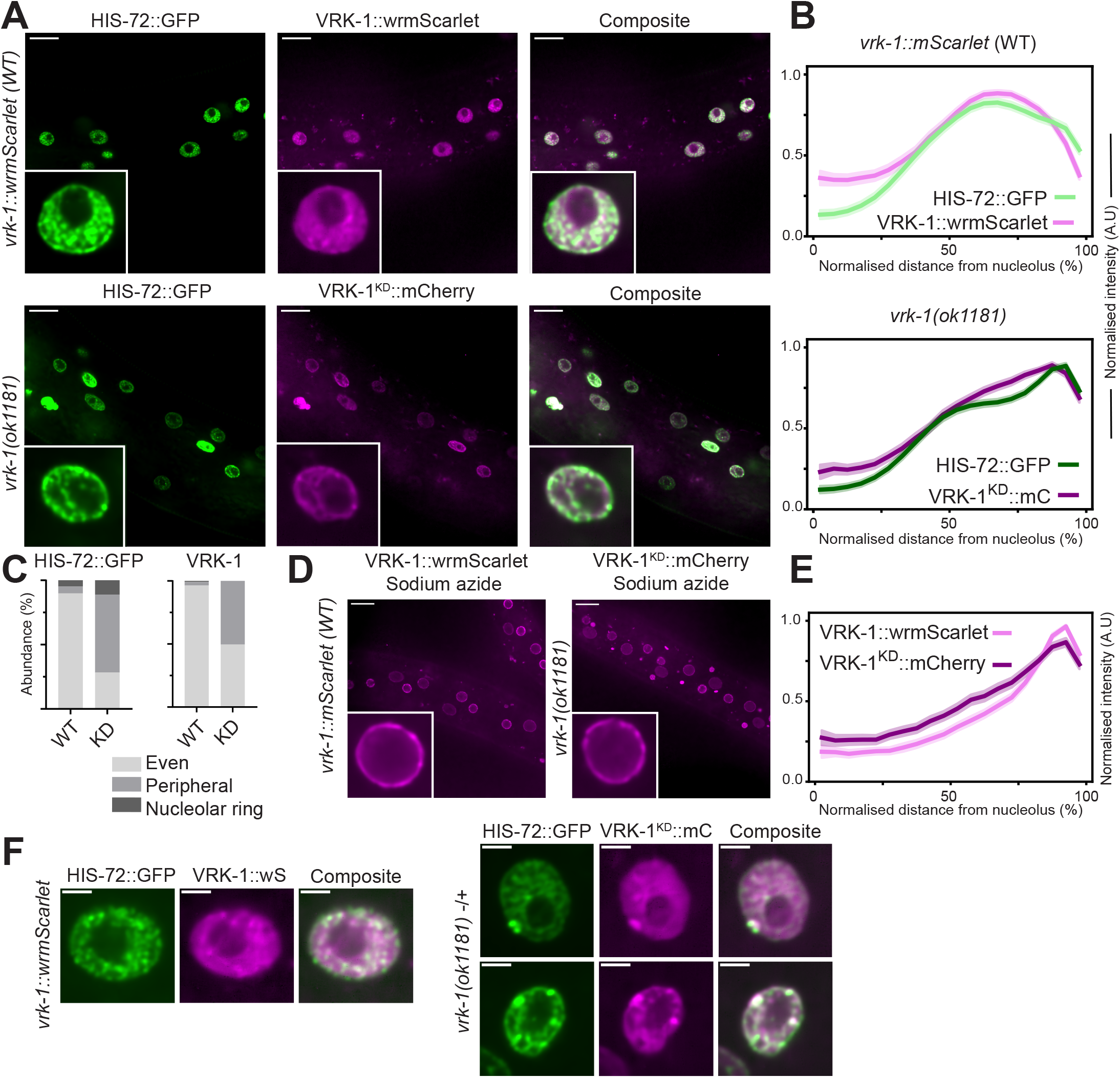
Catalytically inactive VRK-1 results in increased peripheral chromatin in non-stress conditions. **A.** Confocal, single plane, denoised representative images of HIS-72::GFP/VRK-1::wrmScarlet (top) and HIS-72::GFP/VRK-1^KD^::mCherry in *vrk-1 (ok1181)* background (bottom) in L4/YA hypodermal nuclei. Scale bar 10 μm. **B.** Average plot profiles for images displayed in A., measured as described prior. (n=352, n=256, respectively) (top) and HIS-72::GFP/VRK-1^KD^::mCherry in *vrk-1 (ok1181)* background (n=459, n=222, respectively) (bottom) of nuclear distribution as described in 1E. **C.** Nuclear categorization from quantifications, measured as described in Fig. 3DE. **D.** Representative single plane images of VRK-1::wrmScarlet (top) and VRK-1 (K169E)::mCherry (VRK-1^KD^::mCherry) in hypodermal nuclei (bottom) after being immobilised with 0.1% sodium azide agarose pads for 20 minutes, as described in Fig. 2. **E.** Average plot profiles for VRK-1::wrmScarlet or VRK-1^KD^::mCherry on sodium azide, as described prior. (n=219, n=230 for VRK-1::wrmScarlet and VRK-1KD::mCherry, respectively). Scale bar 10 μm. **F.** Hypodermal nuclei in young adults expressing HIS-72::GFP and VRK-1 mScarlet (top) and HIS-72::GFP/VRK-1^KD^::mCherry in *vrk-1 (ok1181)* heterozygotes. Scale bar 2 μm.

Furthermore, VRK-1^KD^ itself was localized at the nuclear periphery, largely overlapping with chromatin distribution, suggesting it retains its ability to bind to chromatin (Fig. 5A–C, S5A). Previous reports suggest that mammalian VRK1 can activate itself through autophosphorylation in response to various input signals (Yokobori et al. 2020; Lopez-Borges and Lazo 2000; Lazo 2024)). We therefore measured whether the relocation of VRK-1 to the nuclear periphery after sodium azide treatment was dependent upon its catalytic activity. VRK-1^KD^ largely maintained the ability to relocate to the nuclear periphery upon azide treatment, albeit with a mild reduction in peripheral enrichment compared with wild-type (Fig. 5DE, S5B). This could indicate that regulatory inputs other than VRK-1 itself govern sodium azide-induced relocation, or that redundant factors act alongside VRK-1 to drive its relocation.

Interestingly, in *vrk-1(ok1181)* heterozygotes (therefore with wild-type VRK-1), we frequently observed large domains of compacted chromatin enriched for VRK-1^KD^, which we never observed for wild-type VRK-1 (Fig. 5F). This suggests that catalytically inactive VRK-1 retains chromatin-binding capacity but fails to dissociate from compact chromatin, possibly outcompeting wild-type VRK-1, which in turn leads to increased chromatin compaction.

Together, these data demonstrate that VRK-1 kinase activity is required to prevent chromatin from becoming compact and accumulating at the nuclear periphery, but is dispensable for overall VRK-1 localisation to chromatin and relocation to the NE on sodium azide.

### BAF-1 is the major target of VRK-1 to prevent overretention of chromatin at the nuclear periphery

We next investigated the downstream targets mediating VRK-1’s chromatin-regulatory function. Overwhelming evidence suggests VRK1/VRK-1 is the major kinase responsible for phosphorylating BANF1/BAF-1 (Nichols et al. 2006; Lancaster et al. 2007; Bar et al. 2014; Molitor and Traktman 2014; Marcelot et al. 2021; Tang et al. 2023; Paouneskou et al. 2025). In mammalian cells, VRK1-mediated phosphorylation of BAF1 reduces its affinity for chromatin and nuclear envelope components (Lancaster et al. 2007; Molitor and Traktman 2014; Marcelot et al. 2021). Moreover, it has recently been demonstrated during meiosis in *C. elegans*, VRK-1 regulates chromosome detachment from the nuclear periphery via phosphorylation of BAF-1 (Paouneskou et al. 2025). We therefore tested whether depleting BAF-1 could suppress the perinuclear chromatin retention observed in VRK-1^KD^ animal somatic hypodermal cells. RNAi-mediated *baf-1* knockdown in wild-type VRK-1 animals had no detectable effect on chromatin distribution (Fig. 6A–C, Fig. S6A). In contrast, *baf-1*(RNAi) dramatically reduced perinuclear chromatin accumulation in VRK-1^KD^ animals: the proportion of nuclei with peripheral and/or nucleolar VRK-1^KD^ enrichment fell from 71% to 28% (Fig. 6A–C, S6A).

**Figure 6.**
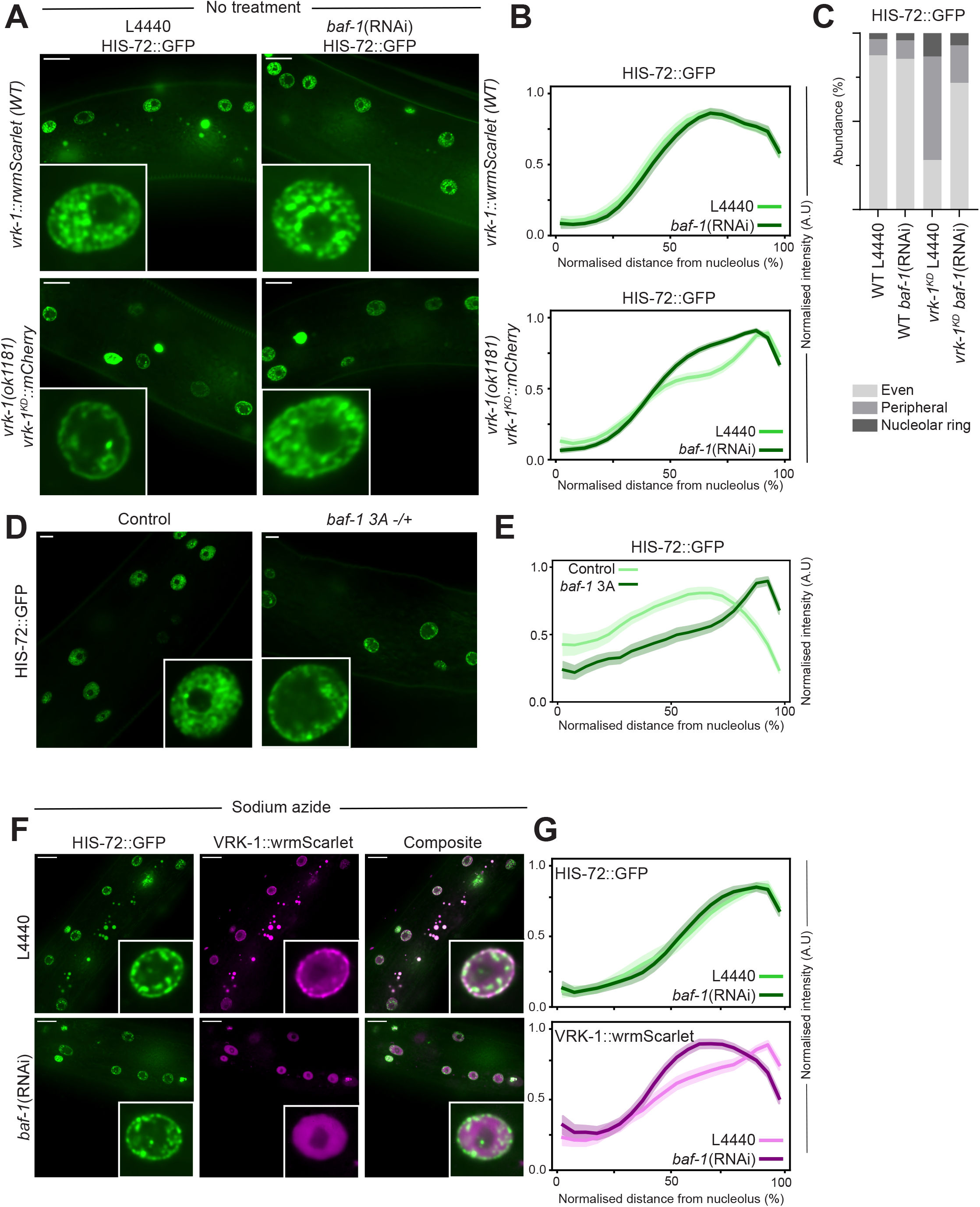
BAF-1 is the major target of VRK-1 to prevent over retention of chromatin at the nuclear periphery. **A.** Hypodermal HIS-72::GFP signal in *vrk-1::wrmScarlet* (top) or *vrk-1p::vrk-1*^KD^*::mCherry*; *vrk-1 (ok1181)* background (bottom). Adult progeny of L4 animals were imaged, grown on *E. coli* expressing either a control empty vector (L4440) (left) or an RNAi construct targeting *baf-1* (right). Scale bar indicates 10 μm. **B.** Average plot profiles for quantifications of images shown in B. (n=143, n=123 for L4440 *vrk-1::wrmScarlet WT, vrk-1*^KD^*::mCherry*, respectively. n=378, n=447 for *baf-1* (RNAi), *vrk-1::wrmScarlet WT, vrk-1*^KD^*::mCherry*, respectively). **C.** Bar charts displaying relative abundance of HIS-72::GFP nuclear distribution categories, measured as described in Fig. 3DE. **D.** Representative images of HIS-72::GFP in wildtype *baf-1* or heterozygous for the unphosphorylatable *baf-1* 3A mutation (see supplementary figure 6 and methods). **E.** Average radial plot profiles for quantifications of D. (n>235 for WT *baf-1* or *baf-1* 3A). **F.** Representative images in adult hypodermal nuclei of HIS-72::GFP and VRK-1::wrmScarlet when immobilised on 0.1% sodium azide agarose pads for 20 minutes with empty vector control (L4440) (top) and *baf-1* (RNAi) (bottom). Scale bar 10 μm. The RNAi treatment was performed as described in A. **G.** Average radial plot profiles for quantifications for F. (n=136, n=113 for HIS-72::GFP L4440, HIS-72::GFP *baf-1* (RNAi), respectively; n=135, n=118 for VRK-1::wrmScarlet L4440, VRK-1::wrmScarlet *baf-1* (RNAi), respectively.

Phosphorylation of BAF-1 by VRK-1 occurs mainly at two highly conserved N-terminal residues, Threonine 3 (T3) and Serine 4 (S4), and possibly Serine 2(S2), with S4 thought to be the most critical phosphosite (Marcelot et al. 2021; Molitor and Traktman 2014). We therefore assessed whether non-phosphorylatable substitutions at these sites would phenocopy loss of VRK-1 catalytic activity. Using CRISPR-mediated genome engineering, we replaced S2, T3 and S4 with alanine residues (*baf-1* 3A; Fig. S6B, see methods). In VRK-1 3A heterozygote mutants, chromatin accumulated at the nuclear periphery, similar to loss of VRK-1 activity, indicating that non-phosphorylatable BAF-1 from at least one allele phenocopies loss of VRK-1 catalytic activity (Fig. 6DE, S6C).

We further examined whether VRK-1’s stress-induced relocation to the nuclear periphery was dependent on BAF-1 by knocking-down *baf-1* using RNAi. Upon sodium azide treatment, *baf-1*(RNAi) abolished VRK-1 perinuclear enrichment while chromatin accumulated at the nuclear rim (Fig. 6FG. S6G). These findings demonstrate that stress-driven chromatin compaction and perinuclear enrichment are BAF-1-independent, whereas VRK-1’s targeting to the nuclear periphery requires BAF-1. Importantly, together these data position BAF-1 as the critical VRK-1 substrate mediating chromatin release from, and/or retention at the nuclear envelope.

### VRK-1 depletion leads to widespread decrease of phosphorylation during heat-shock recovery

As a multifunctional nuclear kinase (Lazo 2024), VRK-1 likely acts on nuclear substrates beyond BAF-1. To identify additional targets, we performed phosphoproteomics comparing VRK-1-depleted and control animals 24 h after heat shock where chromatin remains compacted in VRK-1-depleted animals but has recovered in controls (Fig. 3).

We identified 29,310 unique phosphosites, of which 1142 showed decreased phosphorylation (mapping to 491 proteins) and only 173 showed increased phosphorylation, representing 107 proteins (Fig. 7A; Supplementary Table 4). Among the proteins with decreased phosphorylation, 27 overlapped with the H3K9me3 ChromID proteome, including the transcription factor LIN-15b, histone H3 (HIS-71/72), Y77E11A.7, and SET-26. The phosphorylation sites on HIS-71/72 (S58, T59) are adjacent to H3K56me3, a histone mark that partially colocalizes with H3K9me3 genome-wide (Jack et al. 2013). No BAF-1 peptides were detected, likely owing to the small size of BAF-1 (89 amino acids), which yields tryptic peptides below the detection threshold.

**Figure 7.**
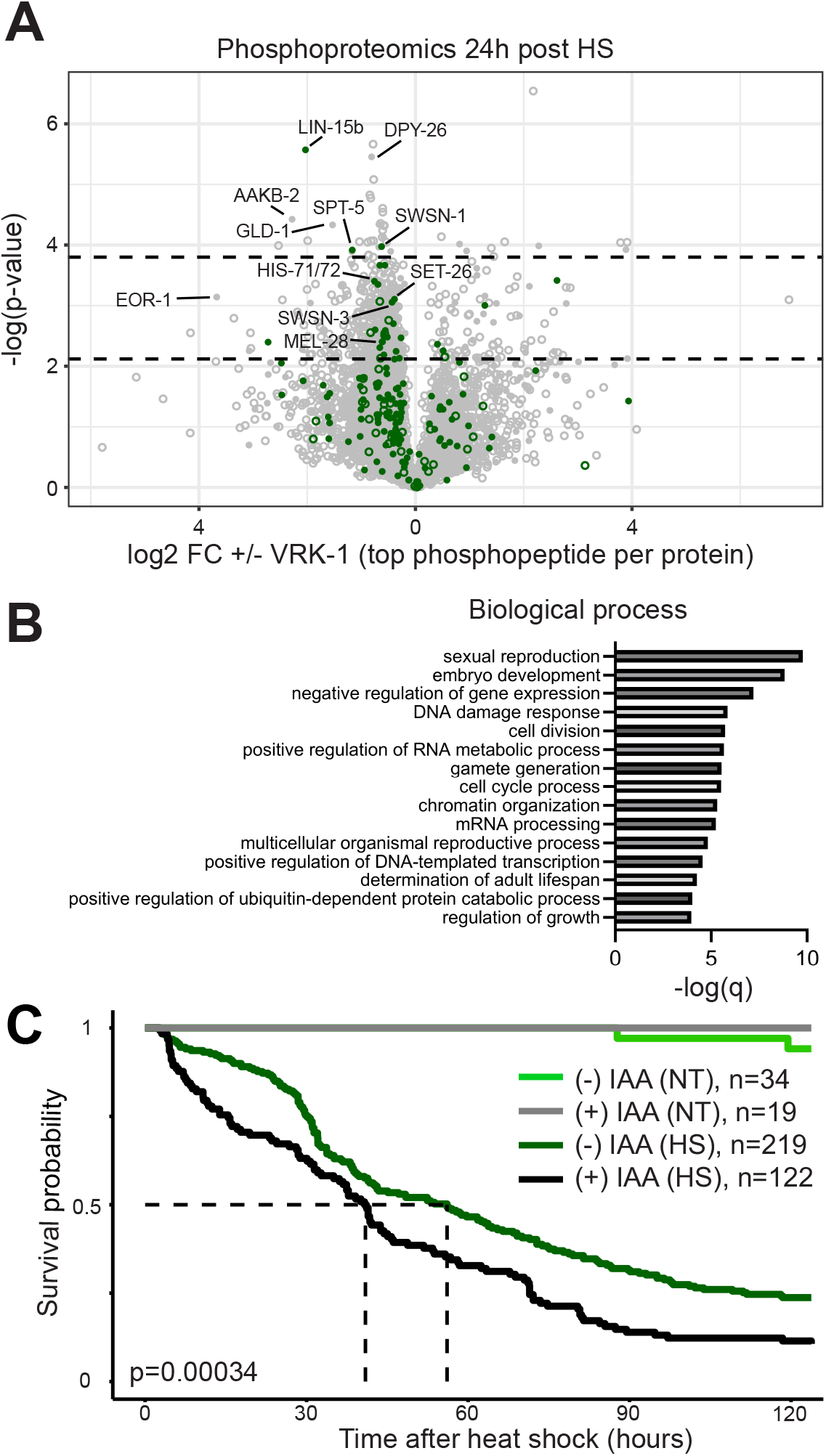
Depletion of VRK-1 affects the phosphoproteome 24h post heat shock and lowers survival after heat shock. **A.** Label-free phosphoproteomics. Volcano plot of differentially enriched phosphopeptides 24 post heat-shock with or without IAA, with the most significant peptide displayed for each protein. The fold changes are log2 transformed. Two statistical cutoff points were used (-log (p)>2.12, low confidence hits), (-log (p)>3.8, high confidence), corresponding to 0.05≤q<0.1 and q<0.05 respectively. Approximately 50,000 animals were grown per sample for each condition in five biological replicates. The third replicate was censored from analyses after hierarchical clustering identified the control sample as an outlier. **B.** Gene enrichment analysis of significantly less phosphorylated proteins after filtering for nuclear predicted or unannotated using cellular component ontology. Non-specific (>500 genes associated with term) and redundant terms were filtered out. **C.** Kaplan-Meier plot showing survival after 90 minute heat shock at 37°C or non-heat shocked controls over a period of 120h in control animals (-IAA) and upon VRK-1 degron-induced degradation (+IAA). Experiment was performed with individual worms in microchambers, heat shock was induced at the third larval stage, and images were acquired every 20 minutes. The number of worms assessed is indicated in the figure. Animal death was manually scored from randomised, blinded, individual timelapse videos.

GO enrichment analysis of nuclear or unannotated proteins with decreased phosphorylation highlighted terms including negative regulation of gene expression, DNA damage response, cell-cycle process, and determination of adult lifespan (Fig. 7B) - functions previously linked to VRK-1 (Campillo-Marcos et al. 2021; Lazo 2024). These results reveal that VRK-1 loss broadly disrupts the nuclear phosphoproteome, consistent with VRK-1 functioning as a pleiotropic chromatin-associated kinase.

### Depletion of VRK-1 reduces survival after heat-shock

In nematodes, *vrk-1* overexpression increases lifespan, whereas *vrk-1* deletion reduces it (Park et al. 2020). We therefore asked whether auxin-mediated VRK-1 depletion, and the associated failure of chromatin to recover its normal distribution after heat stress, affects organismal fitness. Individual animals were tracked for 120 hours in agarose microchambers to evaluate survival with or without a heat shock. Animals were exposed to a slightly longer heat shock (90’ at 37°Cinstead of 60’ previously), as the temperature equilibration of the chambers required longer and therefore 60’ was not sufficient to induce previously observed lethality phenotypes using this system (data not shown) (Stojanovski et al. 2022). Auxin treatment alone had no significant effect on survival in the absence of heat shock over the time course of the experiment (Fig. 7C). VRK-1 depletion, however, significantly reduced survival during heat-shock recovery: after 120 hours, 24% of control animals remained alive compared with 12% of VRK-1-depleted animals (Fig. 7C). These data indicate that VRK-1 is required for thermal stress tolerance, and associate the failure to restore normal chromatin architecture with reduced organismal survival upon stress recovery.

## Discussion

In this study, we validated proximity labeling for H3K9me3 in *C. elegans* using ChromID, revealing a diverse H3K9me3-proximal proteome. Among the factors identified, we focused on the conserved kinase VRK-1. However, we show that VRK-1 chromatin regulatory roles likely extend beyond H3K9me3 regulation: VRK-1 regulates global chromatin architecture, relocates to the nuclear periphery upon stress, and is crucial for chromatin decompaction and release from the nuclear rim during stress recovery. Loss of VRK-1 catalytic activity caused constitutive chromatin accumulation at the nuclear periphery. This phenotype is reversed by depletion of the VRK-1 substrate BAF-1 and expression of non-phosphorylatable BAF-1 phenocopies excessive chromatin accumulation at the NE observed in catalytically inactive VRK-1. This reveals a conserved VRK-1/BAF-1 signalling axis regulating global chromatin architecture in non-dividing nuclei, adding to previous lines of evidence implicating this pathway in the same regard in dividing cells (Lancaster et al. 2007; Paouneskou et al. 2025).

### ChromID identifies the proteome at H3K9me3

ChromID identified 220 high confidence (q<0.05) and 251 (q<0.1) proteins in total associated with H3K9me3, including well-known H3K9me3-associated factors, such as the heterochromatin protein HPL-2 (Garrigues et al. 2015), the chromodomain protein CEC-3 (Greer et al. 2014), the chromodomain protein and H3K9me3 nuclear envelope anchor CEC-4 (Gonzalez-Sandoval et al. 2015), as well as inner nuclear envelope components (LMN-1, EMR-1, LEM-2 and UNC-84) (Ahringer and Gasser 2018), thereby validating this approach. We also detected nucleolus-associated proteins, consistent with the enrichment of H3K9me3 in this compartment (Bizhanova and Kaufman 2021). The absence of SET-25, LIN-61, and HPL-1 - three factors previously shown to interact with H3K9me3 chromatin - likely reflects technical limitations of mass spectrometry, such as poor ionization or inefficient tryptic digestion, as no peptides were detected for these proteins in any of the pulldowns (Padeken et al. 2021; Garrigues et al. 2015). Factors more closely associated with H3K9me2 (CEC-5, LET-418, LIN-15b) (Hou et al. 2023; Rechtsteiner et al. 2019) were also detected, probably reflecting their spatial proximity to H3K9me3. We additionally identified proteins previously associated with euchromatin (MRG-1, SET-26, BET-1) (Wang et al. 2018; Cabianca et al. 2019; Kim et al. 2021b), which may reflect transient interactions that proximity labeling captures more sensitively than co-IP or ChIP-seq (Jablonska 2024).

Components of four different chromatin remodeling complexes were also enriched at H3K9me3 chromatin: the NuRD chromatin remodeling complex (CHD-3, LET-418, HDA-1), classically associated with gene repression (Passannante et al. 2010); the SynMuv/DRM complex (LIN-35, LIN-9, LIN-36, LIN-54, LIN-15b), which regulates the germline/soma barrier via H3K9me2 (Rechtsteiner et al. 2019; Hoareau et al. 2024); the FACT complex (HMG-4, SPT-16), whose yeast homolog stabilizes histone turnover rate at heterochromatin (Murawska et al. 2021; Suggs et al. 2018); and the SWI/SNF complex (SWSN-1, SWSN-3), which is involved in both gene activation and repression through chromatin remodeling (Large and Mathies 2014).

Novel candidate H3K9me3-associated factors include JHDM-1, whose human ortholog KDM2B demethylates H3K9me3 (Lee et al. 2019; Jiang et al. 2019); HMG-11/12, whose ortholog HMGA2 regulates H3K9me3 distribution and mediates chromatin compaction (Mallik et al. 2018; Kuwayama et al. 2023; Divisato et al. 2022); and the SET-domain protein Y77E11A.7 potentially linked to transgene repression (Vastenhouw et al. 2003; Sun et al. 2011). We emphasize that proximity labeling identifies proteins in the vicinity of H3K9me3 - including freely diffusing nuclear factors - and that confirmation of direct association will require orthogonal validation for individual candidates, such as colocalization with known heterochromatin markers or ChIP-based approaches in future studies. ChromID thus provides a broadly applicable platform for mapping histone-mark-proximal proteomes *in vivo*, with potential adaptation to inducible, tissue-specific, or condition-specific contexts such as aging or stress.

### VRK-1 is a component of the H3K9me3 proteome

Among H3K9me3-proximal factors, we identified VRK-1 (Fig. 1). VRKs are a family of conserved chromatin-binding serine/threonine kinases conserved from invertebrates to mammals (VRK-1 in *C. elegans*, NHK-1 in *Drosophila,* hVRK1/2/3 in *H. sapiens*). They have been implicated in numerous, varied functions, including the DNA damage response, cell cycle progression and chromatin organisation (Lazo 2024). This is reflected in their diverse range of substrates including histone H3, BAF, p53 and coilin (López-Sánchez et al. 2014; Cantarero et al. 2015; Molitor and Traktman 2014). Barrier to autointegration factor (BANF1/BAF-1) is a well characterised substrate of VRK1/VRK-1, playing roles in nuclear envelope reassembly post mitosis and in gene expression regulation (Gorjánácz et al. 2007; Romero-Bueno et al. 2024). Importantly, phosphorylation of BANF1 by VRK1 reduces its affinity for chromatin/DNA and components of the nuclear envelope (Lancaster et al. 2007; Molitor and Traktman 2014; Marcelot et al. 2021). With respect to chromatin specificity, loss or catalytic inhibition of hVRK1 increases repressive histone marks (H3K9me3/H3K27me3) and decreases active marks (H3K9ac/H3K27ac) (Monte-Serrano et al. 2023). Similar effects have been observed in *C. elegans* and zebrafish using electron microscopy, suggesting a conserved role for VRKs in negatively regulating heterochromatin (Gorjánácz et al. 2007; Carrasco Apolinario et al. 2023). Consistent with our finding that VRK-1 is associated with H3K9me3, hVRK1 physically interacts with HP1γ, HMGA2 and the H3K9me3 methyltransferase SETDB1 (Monte-Serrano et al. 2023; Lee et al. 2017). In contrast, studies in *Drosophila* show that NHK-1 is associated with the euchromatic H3K27ac histone mark, trithorax group proteins and CBP, acting to maintain gene activation (Khan et al. 2021; Shaukat et al. 2021). More broadly, hVRK1 interacts with a range of other histone modifying proteins including histone deacetylase HDAC1, acetyl transferases PCAF and TIP60, and histone demethylases KDM3A and KDM4A (Monte-Serrano and Lazo 2023; Monte-Serrano et al. 2023). Recent studies reveal hVRK1 phosphorylates High Mobility Group Protein AT-hook 2 (HMGA2) and Chromodomain-helicase-DNA-binding protein 1-like (CHD1L) to regulate gene expression (Gu et al. 2025; Li et al. 2025).

Although VRK-1 was identified through its proximity to H3K9me3, our data suggest that it functions as a general regulator of chromatin–lamina interactions rather than a heterochromatin-specific factor. VRK-1 distribution in unstressed nuclei does not recapitulate the focal perinuclear pattern characteristic of H3K9me3 (compare Figs. 1b and 2a), and its loss affects bulk chromatin distribution regardless of H3K9me3 status. This is consistent with the known biochemistry of VRK family kinases, whose central substrate BAF-1 bridges chromatin to the nuclear lamina in a manner independent of histone modification state. Hence, the chromatin regulatory roles of VRK-1 appear to go beyond the regulation of H3K9me3-marked genes. Given its ability to directly regulate chromatin-modifying enzymes, chromatin binding proteins and histones, VRK-1 has been postulated to act as a master regulator of chromatin dynamics (Lazo 2024).

### Environmental stress induces reversible chromatin compaction

Our findings demonstrate that azide and heat stress trigger rapid yet reversible genome reorganization in which chromatin condenses and accumulates at the nuclear periphery (Fig. 2, 3). Comparable stress-induced genome reorganizations have been reported in mammalian cells under metabolic stress (Visvanathan et al. 2013); (Kirmes et al. 2015) and in *C. elegans* intestinal nuclei during starvation (Al-Refaie et al. 2024). Both active and passive mechanisms likely contribute to this process. Chromatin polymer models predict that reduced transcription promotes compaction and perinuclear anchoring (Cook and Marenduzzo 2009; Ganai et al. 2014); however inhibition of RNA polymerase II or global reduction of kinase activity necessary for transcription fails to mimic this predicted effect (Visvanathan et al. 2013). The speed of the changes we observe argues for active, regulated processes. Directed subnuclear movements of genomic loci have been documented during gene activation (Chuang et al. 2006; Brickner and Walter 2004; Rohner et al. 2013), and during H3K9me2-mediated repression (See et al. 2020). In contrast to these locus specific movements, our data indicate that the majority of the genome shifts towards the nuclear periphery during stress, suggesting the activation of a genome-wide perinuclear anchoring and/or segregation program. Future work should identify the upstream determinants of this reorganization, including potential signaling pathways - some of which genetically interact with *vrk-1* to maintain organismal fitness (Park et al. 2020).

Regarding the function of genome reorganization, the shared morphologies of the genome in animals stressed differently suggest convergence on a common structural endpoint that may be functionally significant. Sodium azide strongly depletes ATP, suggesting passive condensation (Visvanathan et al. 2013), while heat shock elicits broad transcriptional repression with selective activation of stress-responsive loci (Shalgi et al. 2014). Transient condensation and perinuclear positioning may shield sensitive genomic regions (Stephens 2020), preserve nuclear integrity during stress (Schreiner et al. 2015), or facilitate coordinated transcriptional gene regulation (Fischl et al. 2020).

### VRK-1 rapidly relocates and regulates chromatin release from the nuclear periphery

Our data indicates that VRK-1 relocates to the nuclear periphery in response to stress and is required for the reversal of the perinuclear chromatin accumulation during recovery (Fig. 4). Catalytically inactive VRK-1 causes excessive chromatin clustering at the nuclear periphery, which is suppressed by BAF-1 depletion, a major VRK-1 substrate and nuclear periphery component (Fig. 6). This relationship mirrors the situation in *Drosophila* oocytes, where the VRK-1 homolog NHK-1 phosphorylates BAF-1 to release chromatin and enable karyosome formation (Lancaster et al. 2007), and a recent study demonstrating that VRK-1 drives detachment of meiotic chromosomes from the nuclear periphery through BAF-1 in the *C. elegans* germline (Paouneskou et al. 2025). Together, these findings support a conserved VRK-1/BAF-1 signalling axis that modulates chromatin distribution across multiple cell types and developmental contexts. Further, recent findings suggest BAF regulates the anchoring of centromeric heterochromatin blocks at the NE in a manner dependent upon its phosphorylation state in *Drosophila (Torras-Llort et al. 2026)*.

This perinuclear activity could explain the apparent broad range of functions described for VRK1 in mammalian systems, including transcription, heterochromatin regulation, cell cycle, DNA damage response, p53 phosphorylation, epithelial-mesenchymal transition and cancer progression (for review see (Lazo 2024)).

Previous studies have shown that, during heat stress, BAF-1 becomes immobilized at the nuclear periphery in a manner dependent upon VRK-1, (Bar et al. 2014). Whether this immobilisation contributes to chromatin binding or reflects distinct stress-responsive functions remains unclear. We propose that in the absence of VRK-1, BAF-1 remains under-phosphorylated, aberrantly retaining chromatin at the nuclear periphery. Under normal conditions chromatin accumulates gradually (Figs. 4, 5); when heat shock drives strong perinuclear chromatin enrichment and increases spatial proximity to BAF-1, the absence of VRK-1 enhances this binding, leading to perinuclear chromatin accumulation (Fig. 4). Stress-induced relocalisation of VRK-1 to the nuclear periphery (Fig. 8) may prime chromatin for redistribution to the nuclear interior during recovery by modulating BAF-1 phosphorylation and its capacity to bind chromatin. Although BAF-1 emerges as a major mediator of perinuclear genome retention, our phosphoproteomic analyses (Fig. 7) reveal numerous additional chromatin-bound direct or indirect VRK-1 targets during stress recovery, consistent with recent human VRK1 phosphoproteomic analyses (Navarro-Carrasco et al. 2024a, 2024b).

**Figure 8.**
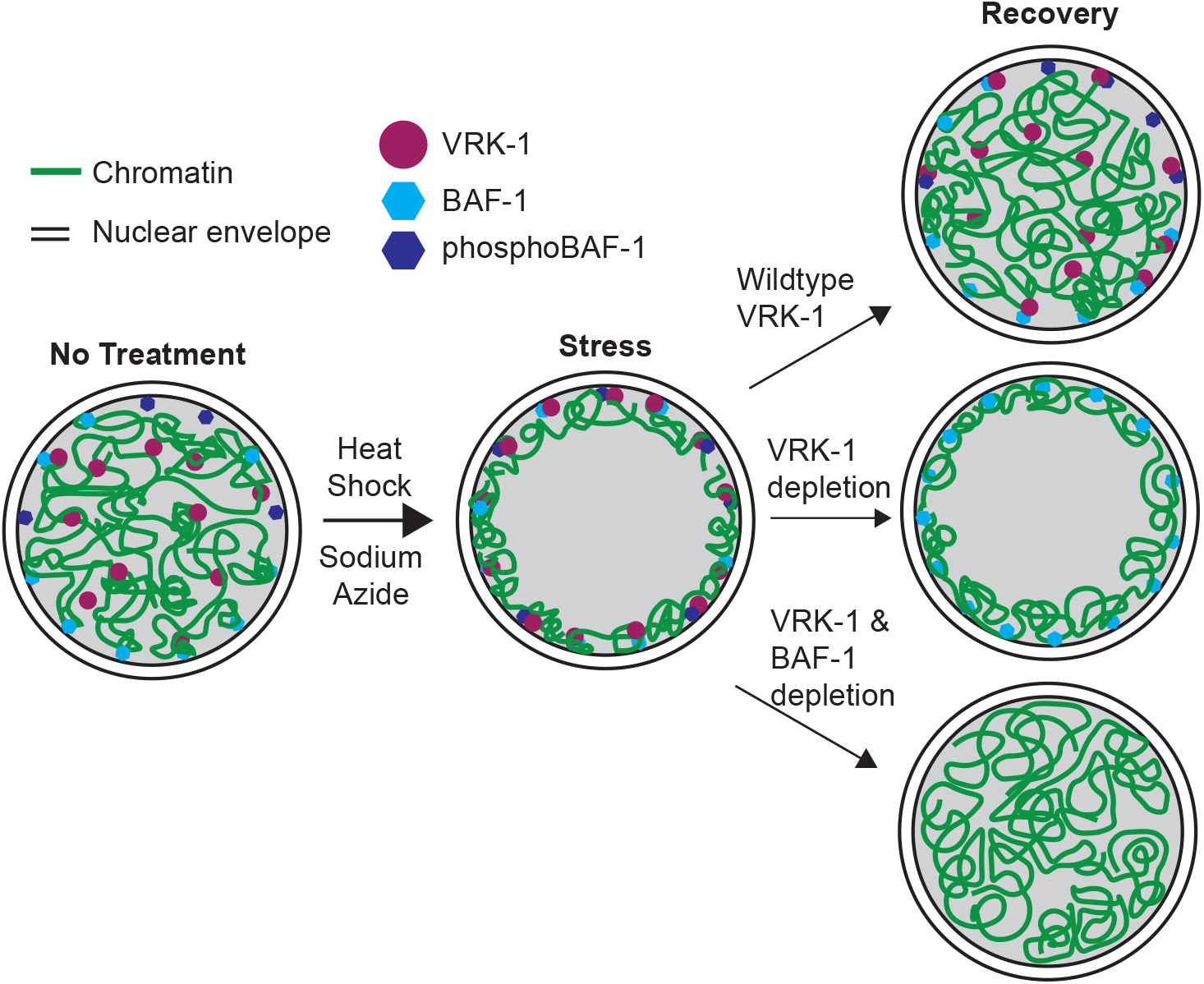
Model where in the absence of VRK-1 chromatin is retained at the nuclear periphery by BAF-1. Green lines indicate chromatin, purple circles indicate VRK-1, light blue hexagons indicate BAF-1 and dark blue hexagons indicate phosphorylated BAF-1. During stress, VRK-1 is targeted to the nuclear periphery followed by compaction of chromatin and its enrichment at the nuclear periphery. In the absence of VRK-1, BAF-1 remains underphosphorylated, thus retaining its ability to interact with chromatin/components of the NE, and ectopically holds chromatin at the nuclear periphery. If BAF-1 is deleted in the background of VRK-1 depletion, chromatin is able to recover its normal distribution.

Together, this study establishes ChromID in nematodes as a platform for mapping chromatin-marks-associated proteomes and identifies the VRK-1/BAF-1 axis that governs chromatin architecture during stress-induced genome reorganization. VRK-1 rapidly relocates to the nuclear periphery during stress and is essential for restoring normal chromatin distribution during recovery. Loss of VRK-1 activity causes persistent perinuclear chromatin compaction even in the absence of stress, underscoring a broader role for the conserved kinase in maintaining chromatin dynamics. Future experiments should determine VRK-1 genome-wide DNA binding pattern, the determinants of its peripheral targeting, and the consequences for gene expression.

## Methods

### *C. elegans* strains and maintenance

*C. elegans* were maintained at 20°C using standard methods unless otherwise stated (Brenner 1974). All strains and genotypes used in this study are listed in supplementary table 1.

### Constructs and strain generation

The CBX1_2x::mScarlet, CBX1_2x::BASU and NLS_BASU were generated by Gibson assembly (see plasmid list) and integrated into a defined site (*ttTi5605*) on chromosome II using MosSCI(Frøkjær-Jensen et al. 2008). Plasmid pBN403 containing wr>mScar>let codon-optimised for *C. elegans* and with two FRT-containing introns (FRT is denoted with a “>” symbol) was generated by Gibson assembly of 3 gBlocks into plasmid pBN312 digested with BsrGI. Next, pBN440 *vrk-1::w>Scar>let* was produced by SapTrap cloning using plasmids pBN403, pMLS256 and pMLS287 and primer pairs B979+B980, B981+B982 and B983+B984.

### Genome engineering

Strain BN1008 *vrk-1(bq28[vrk-1::w>Scar>let])* II was generated by microinjection of plasmid pBN440 *vrk-1::w>Scar>let* at 80 ng/µl into the gonads of HCL67 *uocIs1[eft-3p::Cas9 + unc-119 (+)]* II young adult hermaphrodites together with marker plasmids pRF4 *rol-6(su1006)* (10 ng/µl), pBN40 *lmn-1p::gfp::his-58* (10 ng/µl), pBN41 *myo-2p::gfp* (2.5 ng/µl) and pBN42 *myo-3p::GFP* (5 ng/µl). Worms expressing *vrk-1::w>Scar>let* were crossed first with HT1593 *unc-119(ed3)* III to select second generation Unc progeny (indicative of loss of *uocIs1*) followed by crossing with N2 to restore wild type *unc-119* function. The degron tag was introduced to *vrk-1::wrmScarlet* using CRISPR-Cas9 mediated genomic editing(Paix et al. 2014), with *unc-58* acting as a co-injection marker (see primers, templates and guides for genome editing). BAF-1 3A mutation was generated in the same manner as VRK-1 degron tagging, except *dpy-10* was used as a co-injection marker.

### ChromID sample collection

Mixed stage worms were age-synchronized by hypochlorite treatment, recovering approximately 100,000 animals per sample. Synchronized L1 worms were grown at 22 °C on fresh nematode growth medium (NGM) plates seeded with OP50 bacteria. Worms were harvested at the 22 hour timepoint (L3 stage), washed off the plates with M9 buffer, and incubated in M9 buffer for one hour with washes every 10 minutes to remove free biotin. After the final wash, animals were pelleted, resuspended in 1 ml of M9, combined with one volume of cold, freshly prepared 2x TBS with protease inhibitor mix [1 mM PMSF, 1x E-64, 1x Aprotonin, 1x Pefabloc, 1x Pepstanin-A], flash-frozen as droplets in liquid nitrogen and stored at -80 °C. Samples were collected in triplicate.

### Protein extraction

Frozen worm pellets were cryogenically grinded using a SPEX 6775 cryomill [settings: precool 5 mins, run 1 min, cool 2 min, 2 cycles, 10 CPS]. Ground material thawed on a roller at 4 °C, vortexed and centrifuged for 2 min at 1000 x g. SDS was added to the supernatant to a final concentration of 1%, and the sample was heated at 90°C for 5 min with gentle mixing. After sonication using a Diagenode Bioruptor Standard (10 min, 30 s ON/30 s OFF at 4°C) and equilibration at room temperature, urea was added to 2 M final concentration from a freshly prepared stock solution [8 M urea, 1% SDS, 1x TBS]. After 5 minutes at room temperature, samples were desalted through Zeba Spin Desalting Columns [Fisher] equilibrated with 2M urea, 1x TBS/protease inhibitors. The spin columns were centrifuged for 5 min at 1000 x g. After the first spin, the flowthrough was placed back on the spin column and centrifuged a second time. Protein concentration was measured using a Nanodrop using the Protein A280 setting. This protocol is adapted from (Artan et al. 2022).

### Depletion of mitochondrial carboxylases

His10-tagged carboxylases depletion was performed essentially as described (Artan et al. 2022). Briefly, PureCube Ni-NTA agarose resin (Cube Biotech, 40 um beads) was equilibrated with Ni-NTA binding buffer [100 mM NaH2PO4, 10 mM Tris-HCl, 8 M urea, pH 8.0, 5 mM imidazole] and 3 mg of protein lysate was incubated overnight at 4 °C on a roller. The carboxylase-depleted supernatant was recovered after centrifugation (5 min, 1000 x g) and stored at 4 °C.

### Streptavidin bead preparation and pulldown

Pierce High capacity Streptavidin agarose resin (100 µl, Fisher) was washed three times for 10 minutes with Buffer 1 [50 mM HEPES, pH 7.8, and 0.2% Tween-20] then acetylated with 10 µl of sulfo-NHS-acetate (dissolved in DMSO) for one hour a RT. After three 10 min washes in Buffer 2 [50 mM ammonium bicarbonate, 0.2% Tween-20] and one wash in Buffer 1, prepared beads were combined with carboxylase-depleted lysate and incubated overnight at RT. Beads were washed stringently: two washes with SDS wash buffer [150 mM NaCl, 1 mM EDTA, 2% SDS, 50 mM Tris-HCl, pH 7.4], one wash with TBS-T buffer [150 mM NaCl, 50 mM Tris-HCl, 0.2% Tween-20, pH 7.6], two washes with KCl-T wash buffer [1 M KCl, 1 mM EDTA, 50 mM Tris–HCl, 0.2% Tween-20, pH 7.4], two washes with Na_2_CO_3_-T buffer [0.1 M Na_2_CO_3_, 0.2% Tween-20, pH 11.5], two washes with urea-T buffer [2 M urea, 10 mM Tris-HCl, 0.2% Tween-20, pH 8.0], and five washes with TBS buffer [150 mM NaCl, 50 mM Tris–HCl, pH 7.6]. Bead pellets were stored at -80 °C.

### On-bead trypsin digestion

Bead-bound proteins were denatured in 200 μl denaturation buffer (DB) [8M urea, 100mM Tris-Cl, pH8] with 1 mM DTT for 30 min at RT, filtered to 10kD MWCO filters, alkylated with 5.5 mM iodoacetamide (10 min in the dark at RT with agitation), washed three times with100 mM ammonium bicarbonate in water, pH 8 (ABC), and digested overnight with 3 μg of trypsin diluted to 130 μl ABC buffer at 37°C with agitation. Peptides were eluted by centrifugation, acidified to pH < 2 with 50 % trifluoroacetic acid, and purified using StageTips (Rappsilber et al. 2007).

### ChromID mass spectrometry

LC-MS/MS measurements were performed on a QExactive HFX mass spectrometer coupled to an EasyLC 1000 nanoflow-HPLC (Thermo Scientific). Peptides were separated on self-packed fused silica columns (I.D. 75 μm, ReproSil-Pur 120 C18-AQ, 1.9 μm, 20 cmDr. Maisch) using a gradient of A [0.1% formic acid in water] and B [0.1% formic acid in 80% acetonitrile in water]: samples were loaded with 0% B with a flow rate of 600 nL/min; peptides were separated by 5%–30% B over 85 min at 250 nL/min. Spray voltage was set to 2.3 kV and the ion-transfer tube temperature to 250°C; no sheath and auxiliary gas were used. Mass spectrometry measurements were obtained in data-independent acquisition (DIA) mode. After each survey scan (mass range m/z = 350 – 1,200; resolution: 120,000) followed by 28 DIA scans (30.4 m/z isolation width, precursors from 350-1,200 m/z, AGC target value 3 x 10^6^, resolution to 30,000 and normalized stepped collision energy to 25.5%, 27% and 30%).

### ChromID mass spectrometry data analysis

MS raw files were analyzed using Spectronaut (directDIA+ workflow, standard settings) with a Uniprot full-length *C. elegans* database and common contaminants. Raw abundances extracted were log2 transformed in Perseus; missing values in the BASU control were imputed with downshifted values from the normal distribution. Proteins with at least two valid values in the CBX1_2x condition were tested by two-sample t-test with permutation-based permutation-based FDR-correction. GO terms were extracted from the *C. elegans* UniProtKB database (August 2025) and merged by protein accession. Ribosomal and mitochondrial proteins were filtered out.

### Phosphoproteomics sample preparation

Paired experimental (VRK-1-depleted, +IAA) and control (-IAA) samples were collected in quintuplicate on consecutive days. Approximately 40,000 worms per sample were collected following the heat shock protocol. MS sample preparation was performed as described (Hu et al. 2019): proteins were denatured in 8 M urea, reduced with (1 mM DTT, 30 min at RT), alkylated (5.5 mM iodoacetamide, 30 min at RT in the dark), digested with Lys-C (4 h at RT), and trypsin (urea concentration 1 M, protease:protein, 1:50), acidified with 50% trifluoroacetic acid, and purified by solid phase extraction (HR-X columns, Macherey-Nagel; wash buffer, 0.1% formic acid in deionized water; elution buffer, 80% acetonitrile/0.1% formic acid in deionized water). Eluates were frozen in liquid nitrogen and lyophilized overnight. Lyophilized peptides were resuspended in 80% acetonitrile/0.1% trifluoroacetic acid for phosphopeptide enrichment using Fe (III)-NTA cartridges on a AssayMap Bravo Platform (Agilent)(Post et al. 2017).

### Phosphoproteomics mass spectrometry

LC-MS/MS was performed on an Exploris 480 coupled to an EasyLC 1200 nanoflow-HPLC or Vanquish Neo UHPLC System (all Thermo Scientific), using the same column type and gradient as above. DIA mode: survey scan (m/z = 400 – 1,200; resolution: 120,000) followed by 34 DIA scans (isolation width 24 m/z; AGC target value 300%; resolution 30,000; stepped NCE 25.5%/27%/30%).

### Phosphoproteomics data analysis

Raw files were analyzed with Spectronaut v19 (directDIA+) using a *C. elegans* UniProt database. Phosphopeptide abundances were log2-transformed, normalized to whole-protein levels, and subjected to two-sample t-tests with FDR correction. The third replicate was censored owing to an outlying control sample. GO term enrichment was performed on nuclear or unannotated proteins with decreased phosphorylation. For proteins with multiple phosphopeptides detected, the most enriched (lowest q-value) phosphopeptide was considered.

### Gene enrichment analysis

Gene enrichment analysis was performed using PANTHER 19.0 (https://geneontology.org/)(Thomas et al. 2022) and redundant terms were filtered. Analysis was performed in August 2025 and September 2025 for CBX1_2xChromID and phosphoproteomics, respectively. Input entries were manually filtered to remove duplicates and split entries with multiple protein names. For phosphoproteomics, only hits that were predicted nuclear or non-annotated by CC were used. The top 15 most significantly enriched terms were plotted after filtering for generic terms (>500 genes associated with them) or if a redundant/very similar term was also in the top 15. Unedited gene enrichment tables are provided in supplementary tables 2, 3 and 5.

### Auxin induced degradation (AID) of VRK-1

For all experiments, TIR1 was expressed under the somatic promoter *eft-3p* (see strains list). Indole-3-acetic acid (IAA) (ThermoScientific) was dissolved in ethanol (1 M stock) and added to NGM at 1 mM final concentration; ethanol alone served as vehicle control. Plates were seeded with OP50 and used within two weeks. VRK-1::degron::wrmScarlet degradation was confirmed qualitatively for each experiment, assessed by loss of red nuclear signal.

### RNAi-mediated knockdown

RNAi by feeding was performed using *E. coli* expressing dsRNA targeting *vrk-1* (Vidal Library(Rual et al. 2004)) or *baf-1 (Kamath and Ahringer 2003)*) with L4440 as empty vector control. Bacteria were grown overnight at 37°C on NGM supplemented with ampicillin, and 1 M IPTG (ApolloScientific) was top-spreaded for a final concentration of 4 mM. For *baf-1* RNAi F0 L4s were allowed to lay eggs for one day, and F1 worms were imaged as young adults after three days. For *vrk-1* RNAi, adult worms were transferred to fresh RNAi plates every two days until day 7 of adulthood.

### *baf-1* 3A mutant selection and imaging

CRISPR was performed by gonadal microinjection using *dpy-10* as the co-injection marker (as described above). Non-phosphorylatable BAF-1 3A is expected to lead to defects in cell division, particularly in the actively dividing germline, ultimately leading to animal sterility (Lancaster et al. 2007). Indeed, this was observed in our injections: out of 70 F1 Rol animals singled across 10 injected mother animals (*dpy-10* heterozygotes), 40% of the animals were sterile as day 1 adults, with no visible germline nor embryos laid. This would not be expected to be the case for heterozygous null mutants, which are fertile (*baf-1* nulls can be maintained as balanced strains). To further confirm that sterile animals were indeed *baf-1* 3A heterozygotes, 19 F1 animals (11 sterile, 8 fertile) were tested by amplifying an amplicon centered on the 5’ end of the gene and carrying a restriction digest of this amplicon with *Acc*I (amplicons from wild-type *baf-1* alleles are cut by *Acc*I, amplicons from *baf-1* 3A alleles are not). 11 out of 11 sterile animals were indeed heterozygotes for *baf-1* 3A based on this criteria whereas 8/8 fertile animals were homozygous *baf-1* wild type. For one sterile and one fertile animal, the amplicon was then sequenced (Fig. S6b), the sterile animal indeed carried the heterozygous *baf-1* 3A mutation. We therefore inferred that sterile worms were at least heterozygous for the 3A mutation, while fertile ones carried wild-type *baf-1* alleles. For imaging, F1 sterile or fertile adults were selected from P0 injected mothers.

### Live imaging

1 M levamisole or 10% sodium azide was added to 2% agarose for a final concentration of 10 mM or 0.1%, respectively. Approx. 50 μl of 2% agarose was used for each pad. Animals were imaged in M9 on a Nikon Ti2 Crest spinning-disk confocal microscope. When imaging VRK-1::wrmScarlet and HIS-72::GFP together, single plane images of the hypodermal layer were acquired owing to the rapid photobleaching of VRK-1::wrmScarlet. For HIS-72::GFP alone, z-stacks of the ventral half of the animals were collected. Figures show denoised, contrast-adjusted images, quantifications used unprocessed images.

### Sodium azide exposures

Animals were incubated on 0.1% sodium azide 1% agarose pads at room temperature for indicated time points before imaging. For experiments assessing recovery from sodium azide, animals were instead washed into 500 μl of M9 buffer containing 0.1% sodium azide and incubated for 30 minutes on a rotator. For the during-exposure timepoint, animals were added straight to agarose pads containing 0.1% sodium azide and imaged. For recovery, azide-containing M9 was removed by washing twice and the animals were transferred to seeded NGM plates and allowed to recover for 1 h. Animals were then picked onto agarose pads containing levamisole and subsequently imaged. Combined sodium azide and IAA-mediated VRK-1 depletion experiments were performed as above, except the M9 also contained 1mM IAA (or vehicle control DMSO), and animals were recovered on NGM plates containing IAA.

### Heat shock procedure

For imaging experiments, L3/L4 animals were placed on 60 mM NGM plates and heat-shocked for 1 h in a PCR block set to 37°C. Recovery proceeded at 20°C in an air-cooled incubator. For auxin experiments, animals were synchronised by 2-3 h egg lay, grown for two days at 20°C on NGM plates, then heat-shocked as above. For phosphoproteomics, embryos obtained by hypochlorite bleaching, hatched overnight in M9, and approximately 50,000 L1 larvae were plated on NGM +/- IAA. After 24 h growth at 22°C, animals were heat-shocked for 1.5 h in a 37°C waterbath, with replicates staggered by 1.5 h to ensure uniform temperature. Worms were harvested after 24h recovery at 20°C.

### Quantifications of nuclear fluorescent distribution

Radial fluorescence profiles were measured using Cell-ACDC (Padovani et al. 2022) and a custom python script as described (Al-Refaie et al. 2024). The mid-plane of nuclei in the mid-dorsal or mid-anterior hypodermal layer or intestine was manually segmented in 2D. Contours were calculated from the nucleolar center (or nuclear center when the nucleolus was not visible) to the nuclear periphery. Fluorescence intensity was measured along each contour, normalised to the maximum and binned into 5% distance intervals (20 bins per contour), before averaging intensity *per* bin and *per* nucleus to get individual average-nuclear plot profiles, with each nucleus represented as a row in heatmaps. For nuclear categorisation, each nucleus was classified by visual inspection into one of four categories: uniform distribution throughout the nucleoplasm (even), peripheral enrichment (peripheral), peripheral and around the nucleolus (nucleolar ring), or peripheral + intranucleolar signal (nucleolar diffuse).

### Heat shock survival assay

Individual eggs were deposited in agarose-based chambers (600 × 600 × 20 µm; 4.5% agarose dissolved in S-basal) as described (Stojanovski et al. 2022) and imaged every 20 minutes at 20°C. After 48 hours, chambers were removed from the microscope and heat-shocked for 90 min at 37°C, then returned to the microscope for continued monitoring. OP50-1 bacteria served as food source. IAA (Sigma) was freshly prepared as a 400 x stock in ethanol. Imaging was performed on Squid microscopes (Cephla Inc), equipped with a Blackfly S-CMOS camera (BFS-U3-63S4M-C, Teledyne Vision Solutions) and a 10x / N.A. 0.4 Olympus objective(Li et al. 2020). Worm segmentation used a UNET++ based deep learning model embedded in a custom-made Python script; death times were manually annotated.

## Acknowledgements

We would like to thank Abdulaziz Jaber, Cristina Ayuso and Cihan Elci for technical help, Jennifer Semple and Francesco Padovani (Helmholtz Munich, Germany) for the AC/DC data analysis framework, as well as the Meister laboratory for discussions. We are grateful to Mario de Bono (ISTA Vienna, Austria) for the gift of the tagged carboxylase strains, and Daphne Cabianca (Helmholtz Munich, Germany), Vincent Dion (Cardiff University), Gary Karpen (UC Berkeley), Helder Ferreira (University of St. Andrews) for critical reading of the manuscript. Some data were acquired on machines supported by the Microscopy Imaging Center (MIC) of the University of Bern. Some strains were provided by the CGC, which is funded by NIH Office of Research Infrastructure Programs (P40 OD010440). This work was funded by the Swiss National Science Foundation 31003A_176226/310030_212472 (to PM), IZCOZ0_189884 (to PM and TB), PCEFP3_181204 (to BT), the Spanish Agencia Estatal de Investigación PID2022-137162NB-I00; doi:10.13039/501100011033, to PA), the Uniscientia foundation (to PM), the University of Bern and the Novartis Foundation for Medical/Biological Research (to PM).

## Author contributions

W.S., T. B. and P.M. conceived the study. Y.L., T.S., R.J.C., and V.H.-P. performed the experiments.W.S., V.A., R.P., R.V., D.S.S., M. S., P. L., P.A. performed the investigation. T.B., P.M., B.T., J.D. acquired the funding and resources. P.M. supervised the study. W.S. and P.M. wrote the original draft of the manuscript. All authors reviewed the manuscript.

## Supplementary information

### Strains used in this study

**Supplementary table 1.**
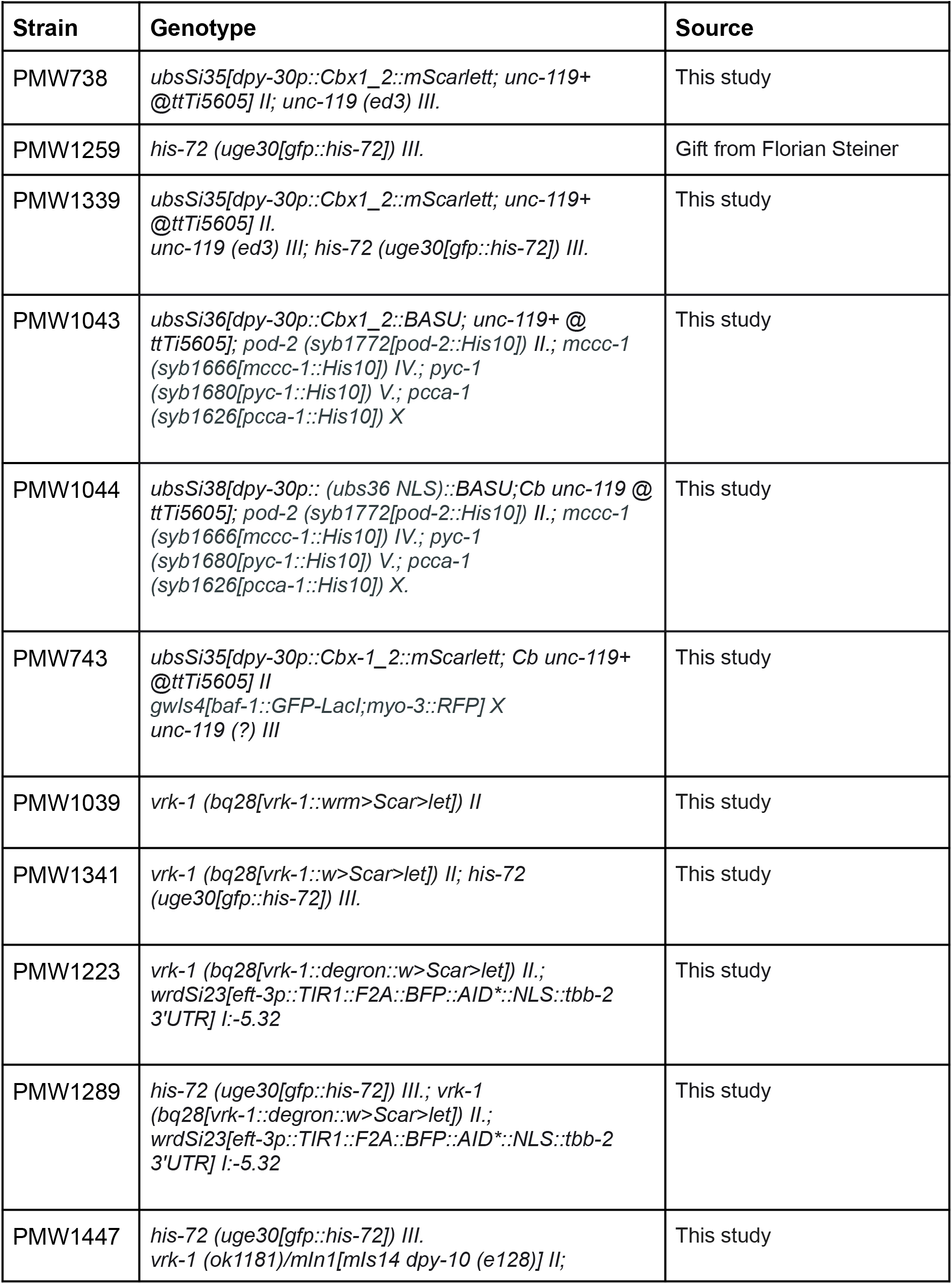
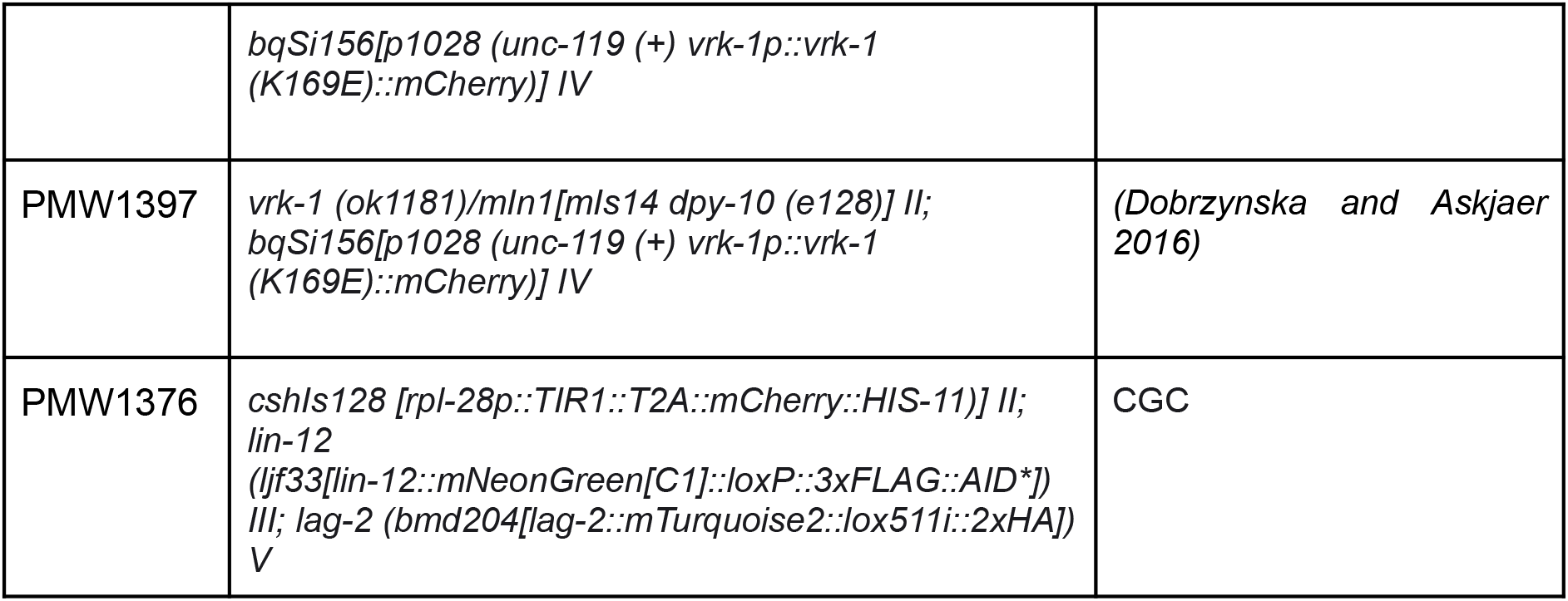
*C. elegans* strains and genotypes used in this study.

### Primers used in this study

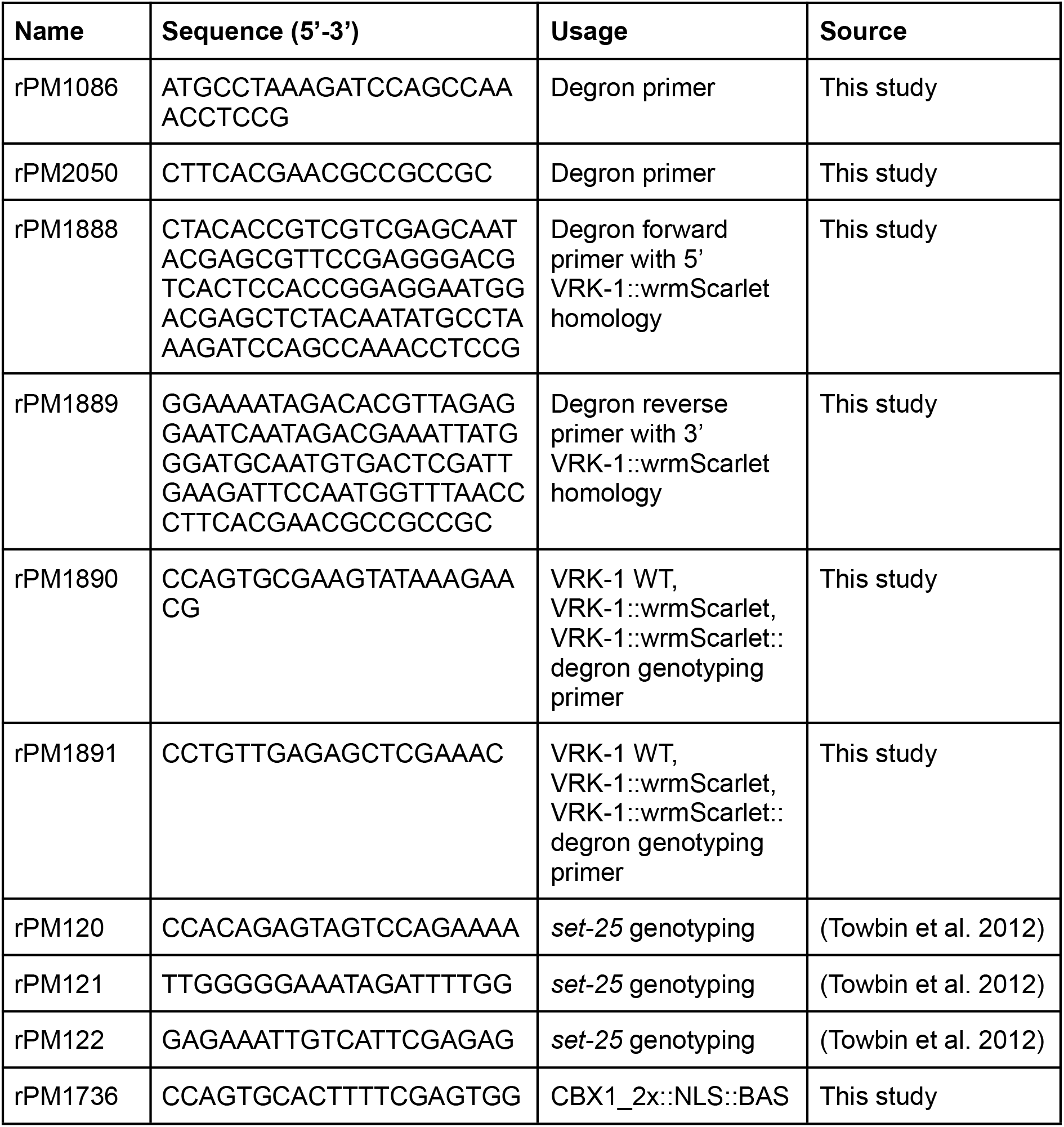
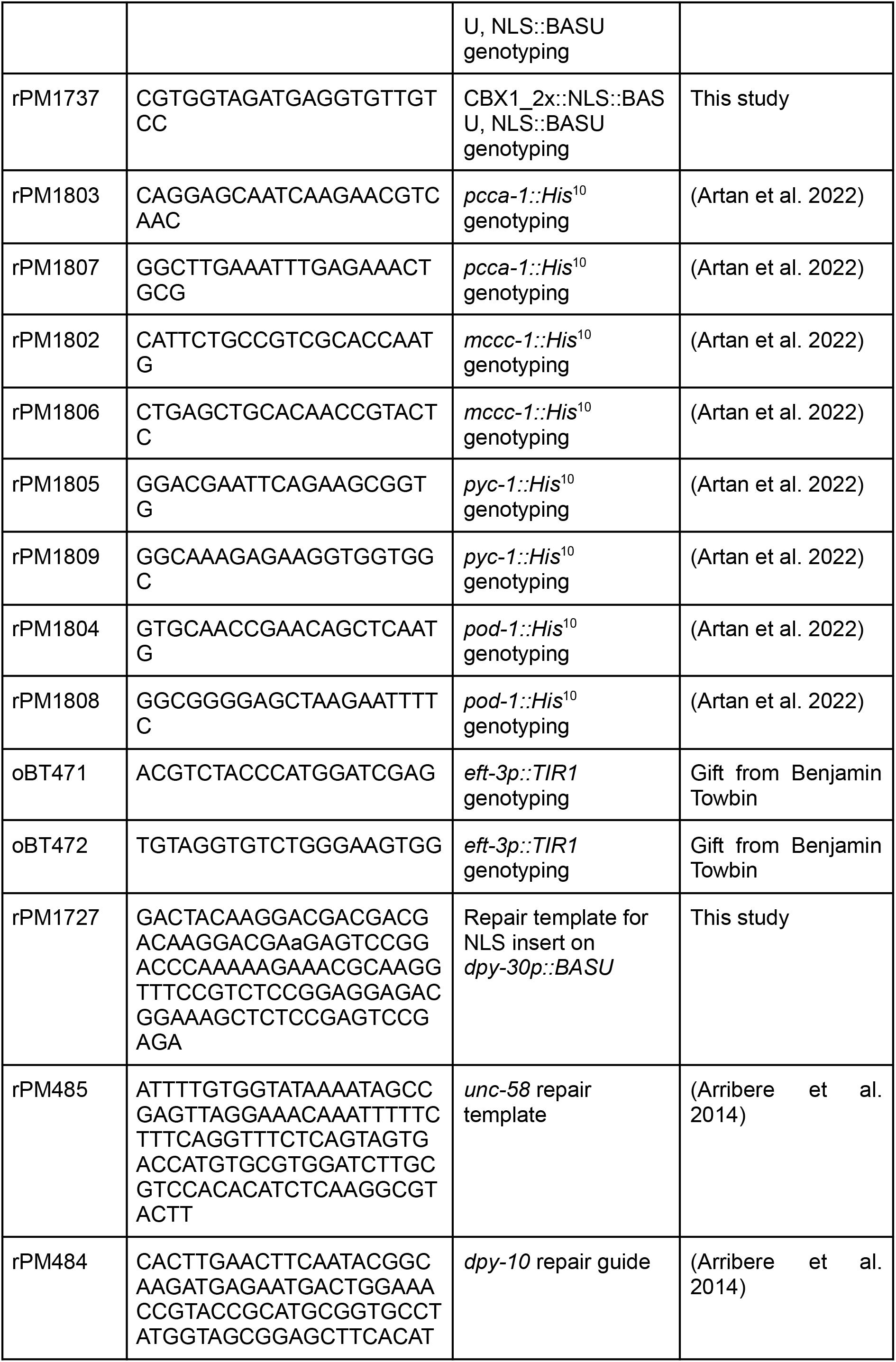
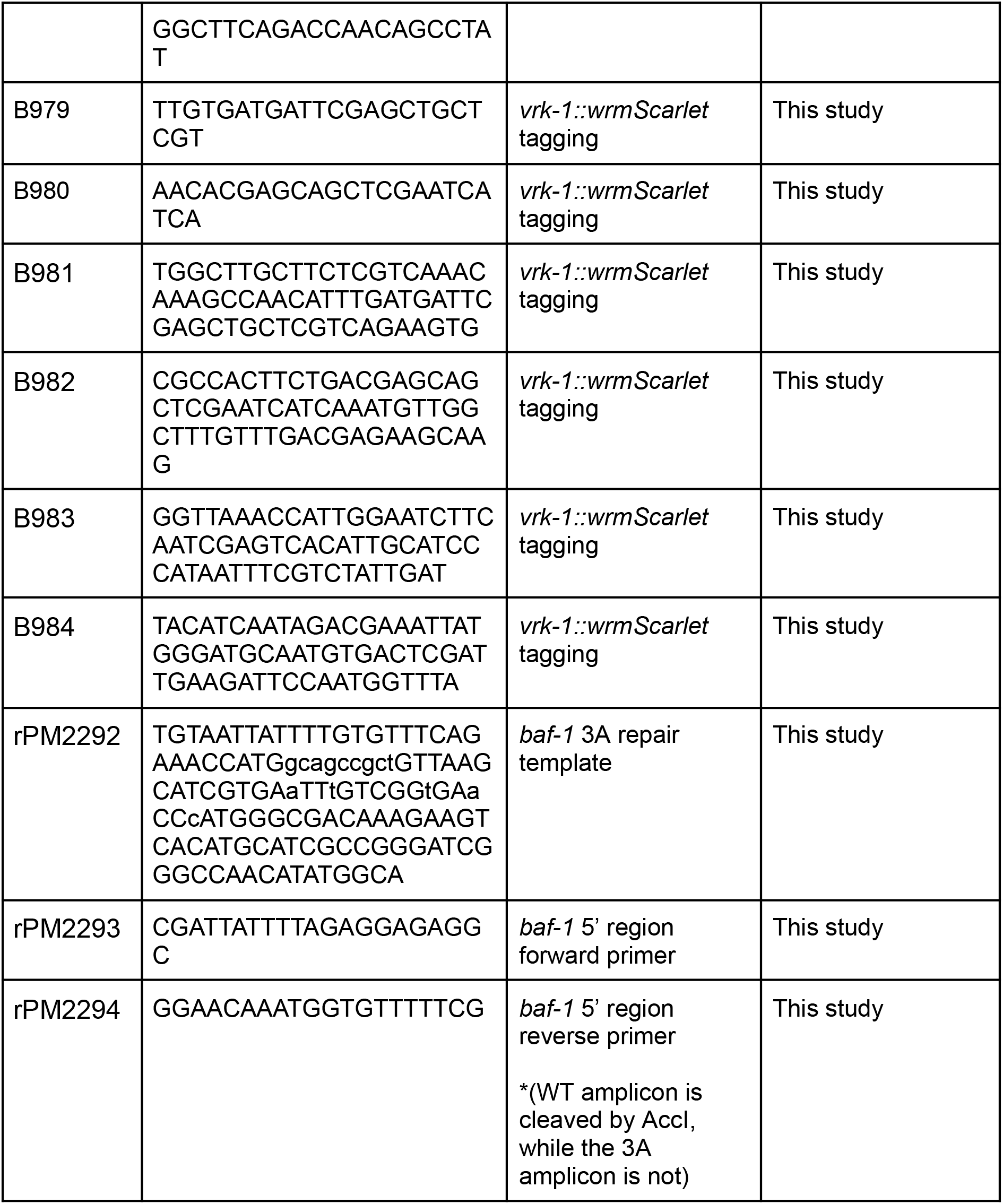

### CRISPR guides used in this study

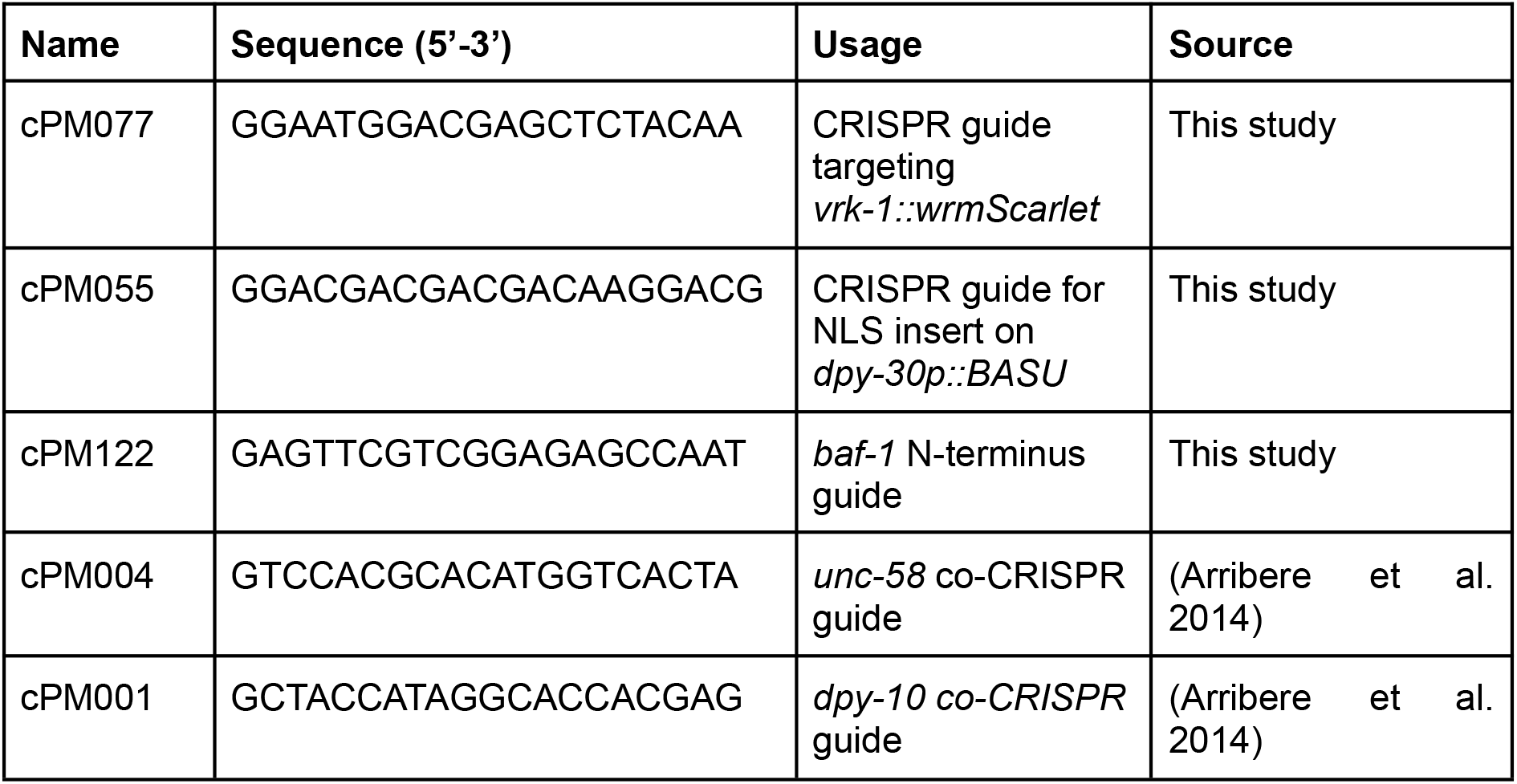

### Plasmids used in this study

| Name | Genotype | Backbone | Source |
| --- | --- | --- | --- |
| pPM495 | *dpy-30p::CBX1_2xChromo::mScarlet::tbb-2 3' UTR* | pCFJ151 | This study |
| pPM497 | *dpy-30p::CBX1_2xChromo::BASU::tbb-2 3' UTR* | pCFJ151 | This study |
| pPM501 | *dpy-30p::BASU::tbb-2 3' UTR* | pCFJ151 | This study |

## Supplementary figures legends

**Supplementary figure 1.**
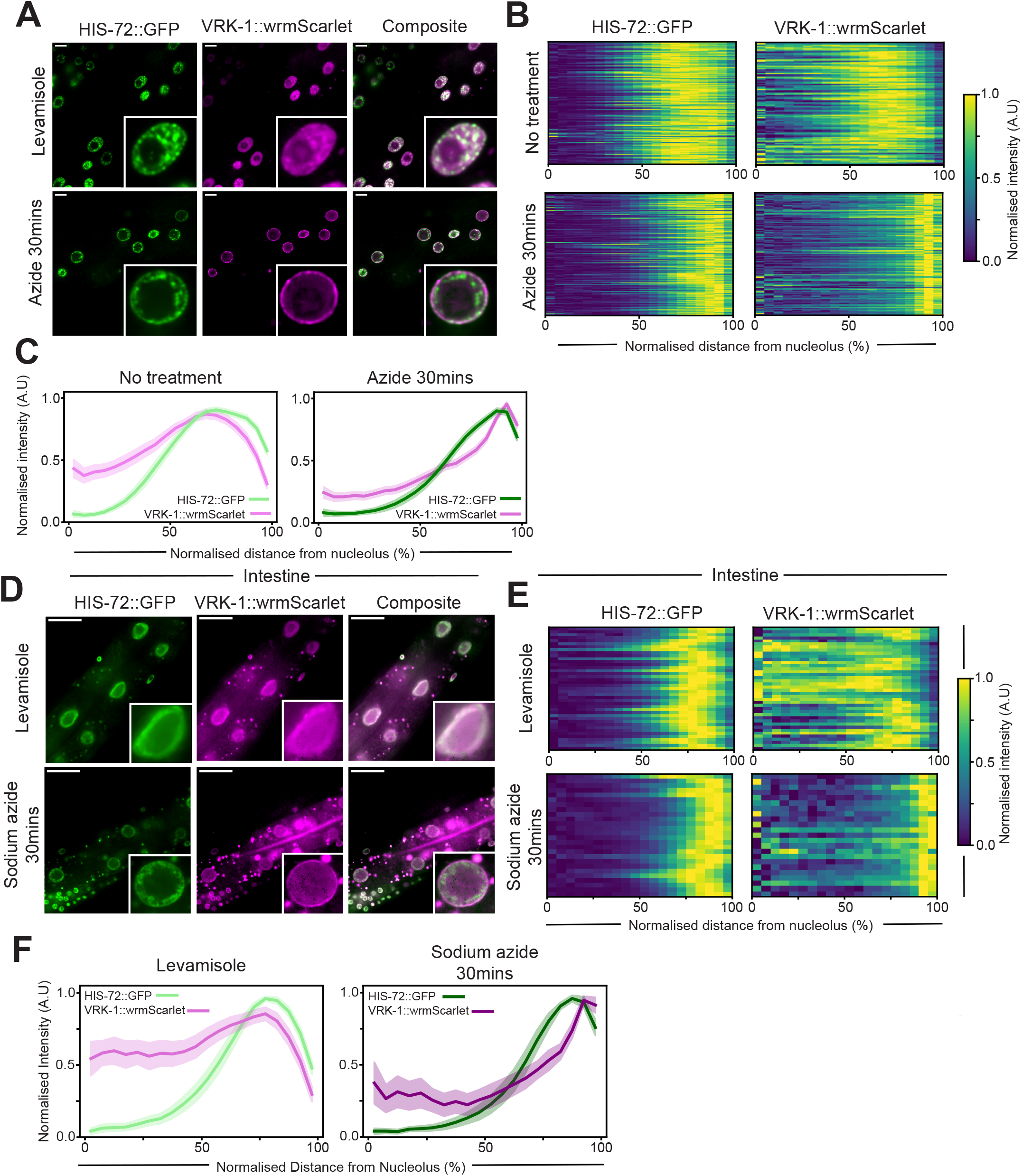
Sodium azide treatment in M9 buffer results in similar VRK-1 and chromatin relocation as agarose pads and VRK-1 relocated to the NE in intestinal nuclei after sodium azide treatment. **A.** Representative confocal single plane images of HIS-72::GFP and VRK-1::wrmScarlet after 30 minute treatment with 0.1% sodium azide in M9 buffer or levamisole, controls for Fig. 2D-F. **B.** Heatmaps displaying normalised intensity from nucleolus/nuclear centre to nuclear periphery for indicated immobiliser or fluorophore, where each row is an individual nucleus for images in A. (n>235 for all conditions). **C.** Average plot profiles for quantifications in B., measured as described in Fig. 1F. **D.** Single plane representative images of intestinal nuclei in L3/L4 larvae on either levamisole or sodium azide as described in A. Scale bar 20 μm. **E.** Heatmaps of normalized fluorescence intensity in intestinal nuclei, as described in B. n=42, n=41 for levamisole and n=27, n=24 for sodium azide 30 mins for HIS-72::GFP, VRK-1::wrmScarlet respectively. **F.** Average plot profiles of intestinal nuclei, as described in C.

**Supplementary figure 2.**
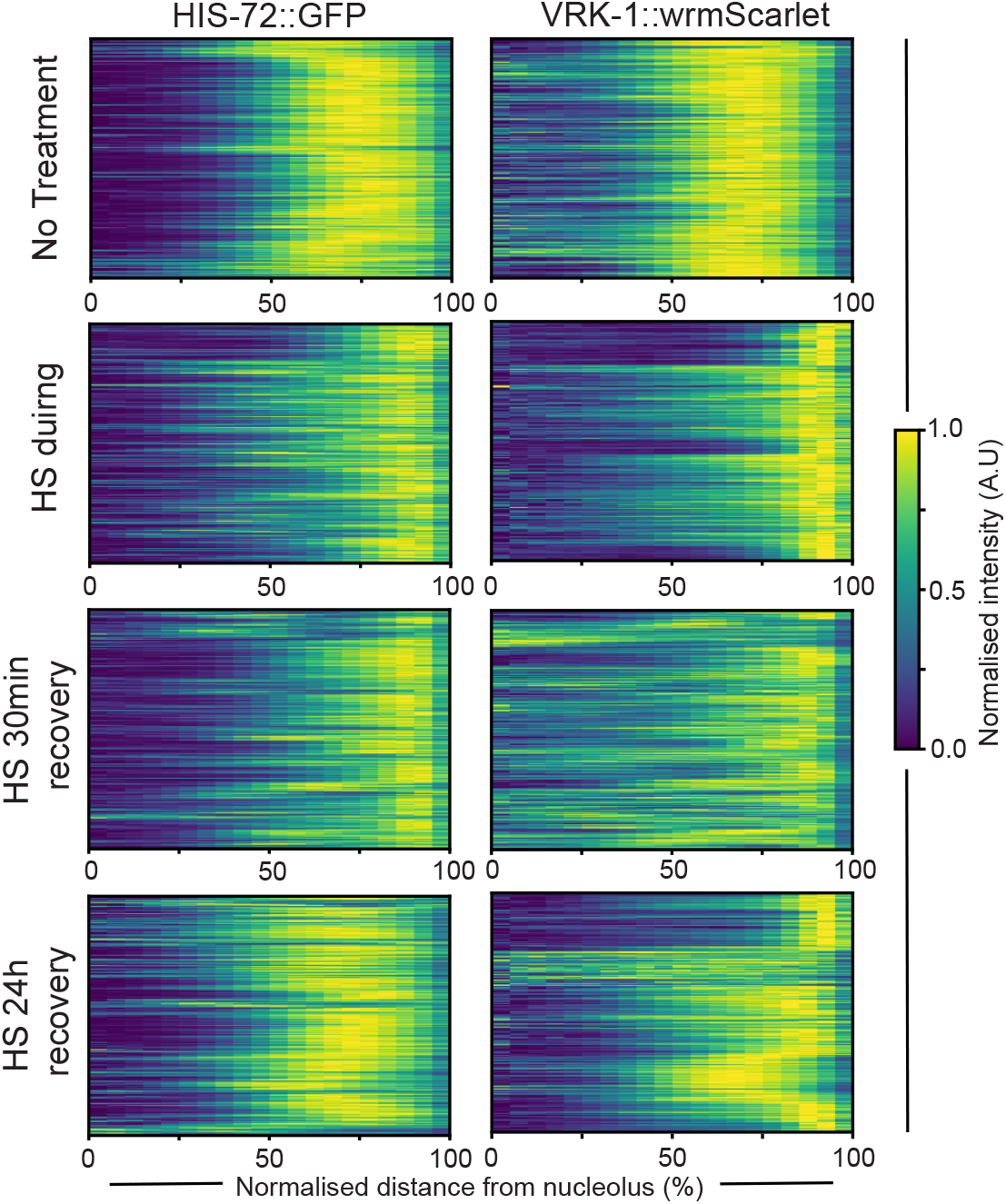
Heatmaps for measurements described in Figure 3. Heatmaps displaying normalised signal intensity from nucleolus/nuclear centre to nuclear periphery for measurements of HIS-72::GFP and VRK-1::wrmScarlet at heat shock timepoints described shown in Fig.3BC. Each row indicates one nucleus.

**Supplementary figure 3.**
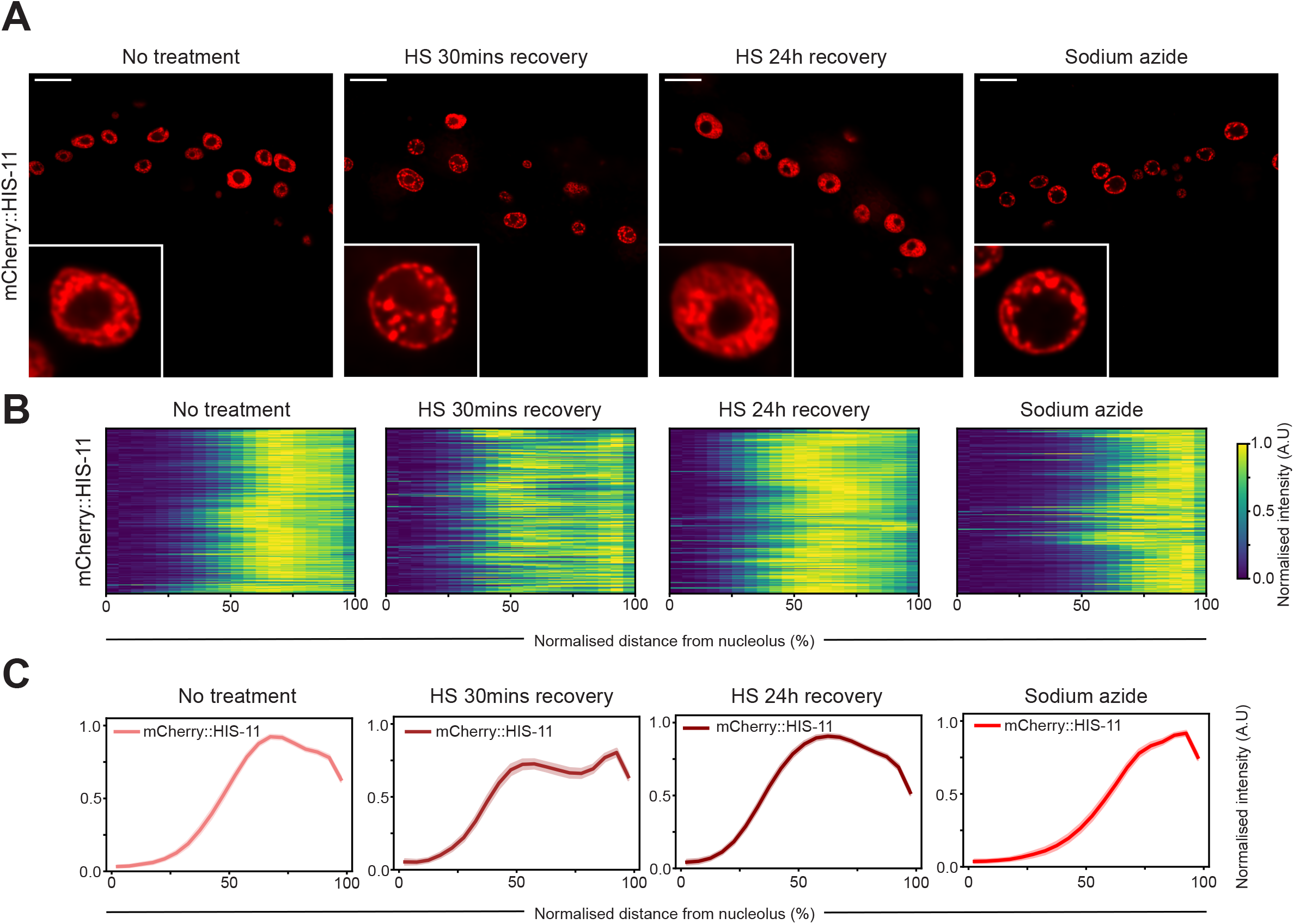
H2B (HIS-11) relocates in a similar manner to H3 in response to sodium azide treatment and heat stress. **A.** Representative single plane images of HIS-11::mCherry in hypodermal nuclei after treatment of levamisole, heat shock (30min recovery and 24h recovery) or sodium azide. Scale bar is equal to 10 μm. **B.** Heatmaps of normalised HIS-11::mCherry distribution from nucleolus/nucleus centre at conditions/timepoints stated in A., where each row corresponds to an individual nucleus. (n>235 for all conditions) **C.** Average plot profiles of HIS-11::mCherry as described in B.

**Supplementary figure 4.**
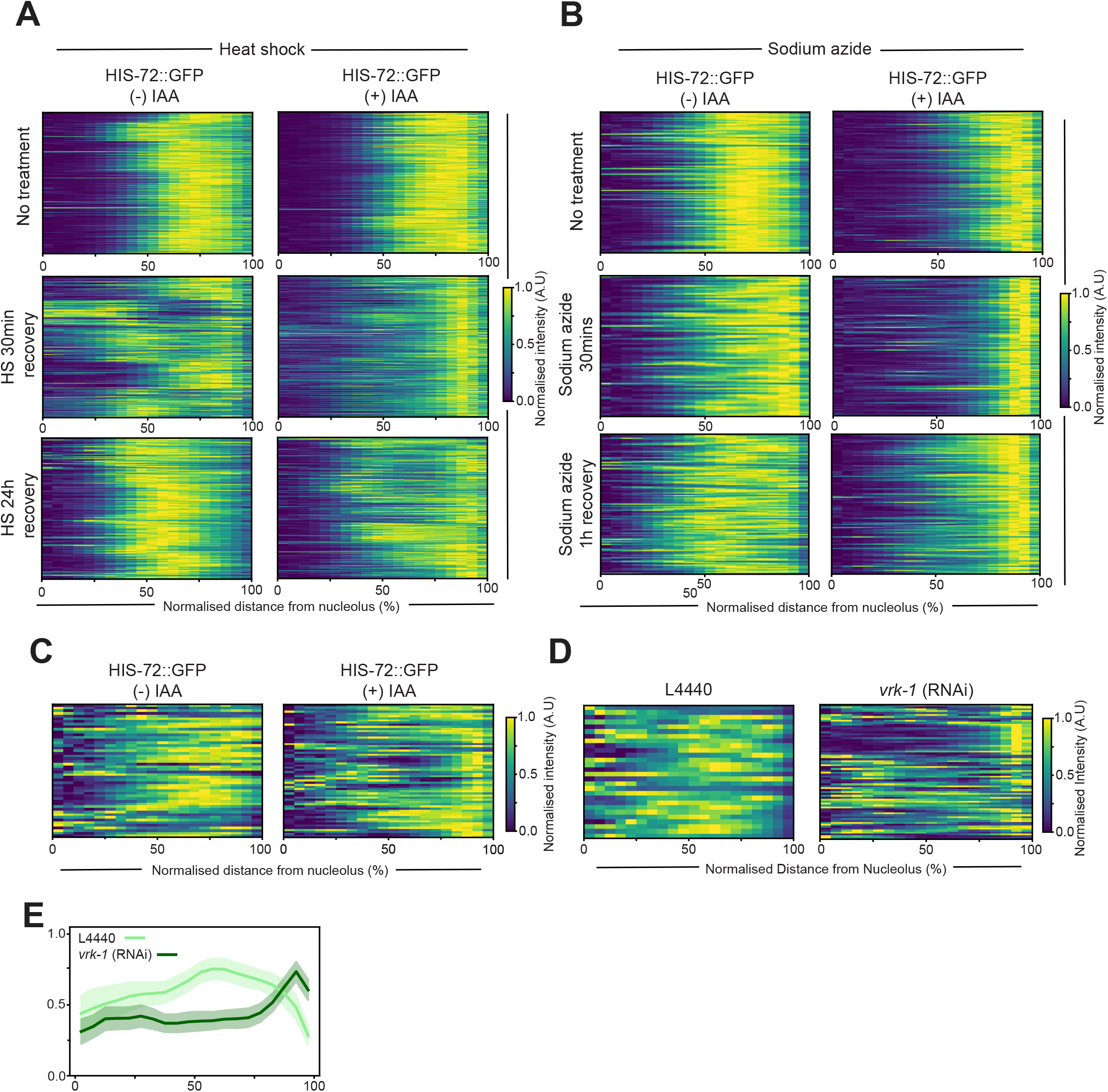
Heatmaps related to figure 4. **A.** Heatmaps displaying quantifications of normalised signal intensity of HIS-72::GFP from nucleolus/nuclear centre to nuclear periphery for Fig.4BC at indicated timepoints before and after heat shock. Nuclear distribution measured as described in 1E. **B.** Heatmaps displaying quantifications of normalised signal intensity of HIS-72::GFP from nucleolus/nuclear centre to nuclear periphery for Fig.4EF at indicated timepoints before and after sodium azide treatment in M9 buffer. **C.** Heatmaps showing quantifications of individual nuclei for day 7 adults, related to Fig. 4GH. **D.** Heatmaps showing quantifications of individual nuclei for day 7 adults after depletion of VRK-1 by RNAi. **E.** Average plot profiles related to D.

**Supplementary figure 5.**
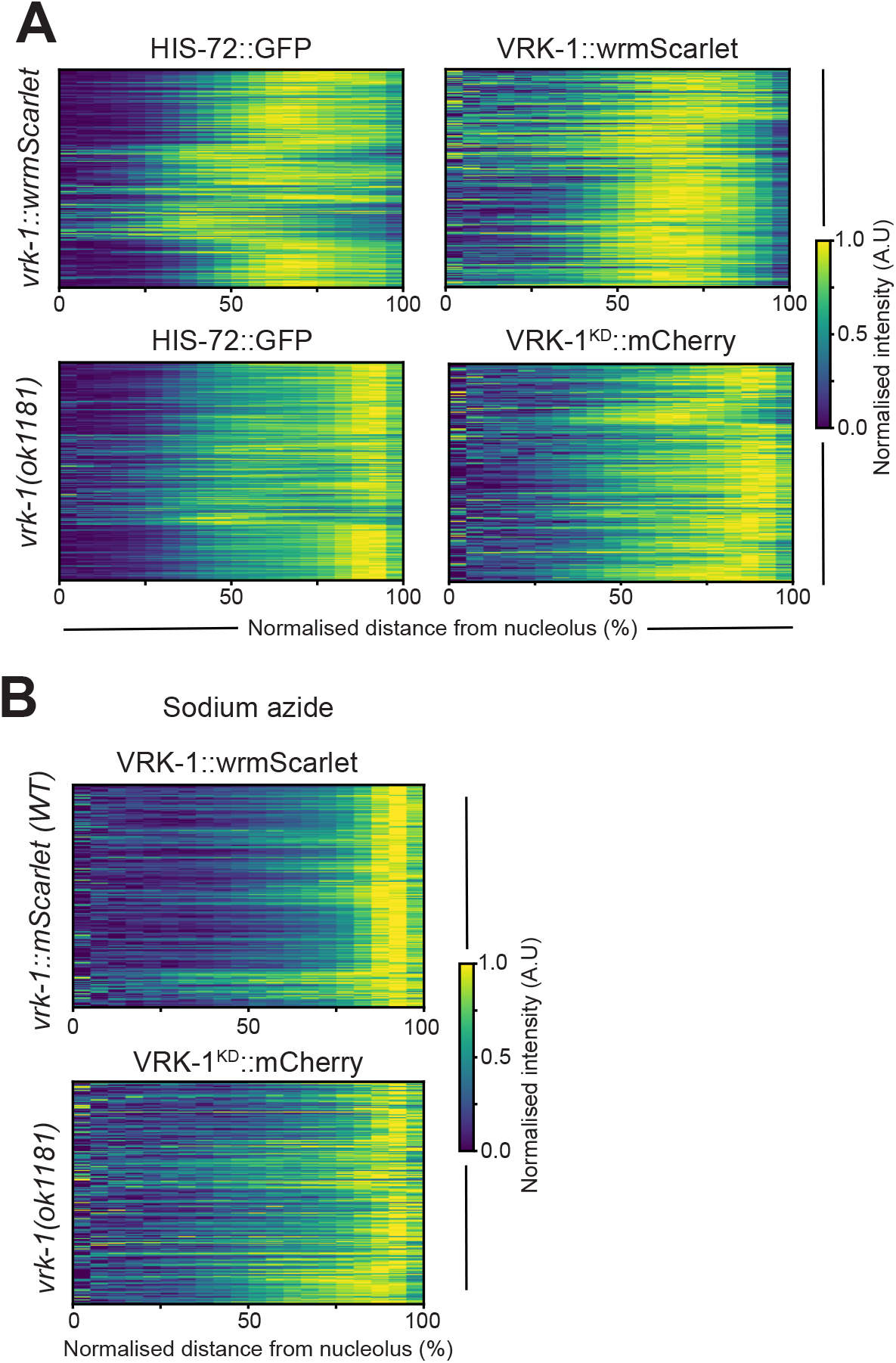
Heatmaps for quantifications from Figure 5. **A.** heatmaps displaying quantifications of HIS-72::GFP/VRK-1::wrmScarlet (n=352, n=256, respectively) (top) and HIS-72::GFP/VRK-1KD::mCherry in *vrk-1(ok1181)* background (n=459, n=222, respectively) (bottom) of nuclear distribution as described in 1E. **B.** Heatmaps of radial fluorescent distribution of nuclei described in A., where each row represents an individual nuclei. n=219, n=230 for VRK-1::wrmScarlet and VRK-1KD::mCherry, respectively).

**Supplementary figure 6.**
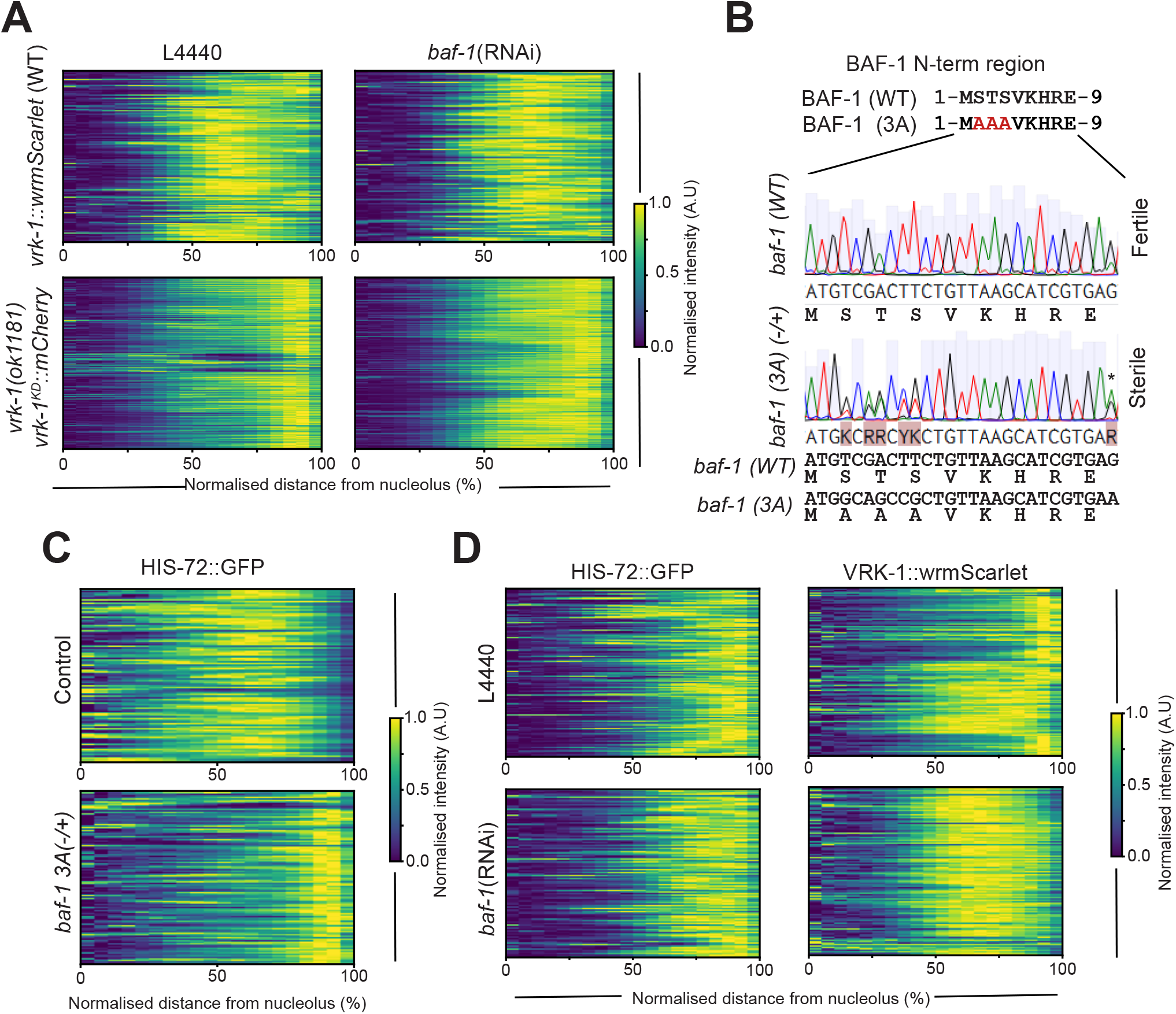
Heatmaps for quantifications related to Figure 6. **A.** Heatmaps showing normalised signal intensity of HIS-72::GFP, where each row indicates an individual nucleus, related to Fig. 6AB. **B.** Schematic showing the substituted amino acids in BAF-1 3A mutant. Sanger sequencing results showing that F1 worms are heterozygote for BAF-1 3A mutation. Note: heterozygotes animals are sterile. Starred mutation at 3’ region is silent, and part of the mutation to ablate the PAM site. **C.** Heatmaps showing normalised signal intensity of HIS-72::GFP, where each row indicates an individual nucleus in WT BAF-1 nuclei (control) and *baf-1* 3A heterozygotes, related to quantification in Fig. 6DE. Genotype was inferred from sterility phenotype, as described in the methods section. **D.** Heatmaps showing normalised signal intensity of HIS-72::GFP, where each row indicates an individual nucleus related to quantifications from Fig. 6FG.

